# *Dexi* disruption depletes gut microbial metabolites and accelerates autoimmune diabetes

**DOI:** 10.1101/393421

**Authors:** LJ Davison, MD Wallace, C Preece, K Hughes, JA Todd, B Davies, CA O’ Callaghan

## Abstract

Non-coding genetic variants in the CLEC16A gene on human chromosome 16p13.13 are associated with risk of autoimmune diseases, including type 1 diabetes and multiple sclerosis. In this region, we previously identified *DEXI*, a candidate causal gene of unknown function, which alters the risk of type 1 diabetes, where the T1D predisposing allele is associated with lower *DEXI* expression. Here, we demonstrate by CRISPR mutagenesis *in vivo* and deep phenotyping that disrupted *Dexi* expression accelerates diabetes in the non-obese diabetic (NOD) mouse, a spontaneous model of autoimmune pancreatic beta-cell destruction. Mutant mice have increased serum IgM and IgA concentrations compared to wild-type NOD mice, as well as changes in both the gut microbiome and molecular metabolites associated with microbial metabolism. These findings suggest that the mechanism by which *DEXI* alters diabetes risk involves the composition and function of the microbiome and its impact on host metabolites. Such metabolites, including short chain fatty acids such as butyrate, have been shown to alter the activity of the immune cells involved in beta-cell destruction and susceptibility of the beta cells to autoimmune attack.

**One Sentence Summary:** Disruption of the *Dexi* gene leads to accelerated diabetes in the non-obese diabetic (NOD) mouse, accompanied by changes in serum immunoglobulins, gut microbiome and microbial metabolites.

## Main Text

### Introduction

Common autoimmune diseases, such as type 1 diabetes (T1D), are caused by a complex, heterogeneous blend of numerous, mostly common, predisposing DNA variants across the genome, alongside a range of environmental factors. Genome-wide association studies (GWAS) have so far mapped over 57 chromosome regions (immunobase.org) affecting T1D risk and, in addition to yielding insights into the inherited mechanisms of disease, also provide clues to identify more elusive environmental factors. Nevertheless for many of these risk regions the actual causal genes and variants are not known. Considerable uncertainty remains around the identity of the environmental factors that, together with a susceptible genetic background, lead to the destructive autoimmune response against the insulin-producing beta cells of the pancreatic islets. Twin, birth cohort and other population studies, along with investigations of animal models, particularly the non-obese diabetic (NOD) mouse, demonstrate the importance of environmental influences in the development of autoimmune disease (*1*), such as exposure to viruses (*2*) and alterations in the gut microbiome (*3*). Some T1D-associated regions identified by GWAS contain genes with known functions in the immune system, several of which are convincing candidate causal genes. Some T1D regions also contain genes with known effects in the target beta cells. In addition to underpinning further understanding of the disease by mapping candidate genes of known functions, GWAS and its follow up studies can identify candidates with no known function at all. Investigation of these genes is challenging, but has the potential to provide entirely novel insights into disease pathogenesis and reveal new therapeutic or preventative strategies.

The T1D-associated 16p13.13 region of the human genome contains multiple genes (**Figure S1**). Single nucleotide polymorphisms (SNPs) in intron 19 of *CLEC16A* in the region have been associated with risk of many autoimmune diseases including T1D (*4, 5*), primary biliary sclerosis (*6*), multiple sclerosis (*7*), primary adrenal insufficiency (*8*), systemic lupus erythematosus (*9*), alopecia areata (*10*) and juvenile arthritis (*11*). Associations with autoimmune disease in number of different SNPs in high linkage disequilibrium have been reported, with the same direction of effect in multiple diseases. A summary of previous association study data is provided in **Supplementary Table S1.** Non-coding T1D-associated variants are in high to moderate linkage disequilibrium with the variants in intron 19 of *CLEC16A* that have been associated with bone density, (*12*), myocardial infarction (*13*), ischemic stroke in metabolic syndrome (*14*) primary IgA deficiency (*15*), combined variable immunodeficiency (*16*), and microbial diversity in the human gut (*17*).

Within the complex 16p13.13 region, most studies have focussed upon the function of *CLEC16A* itself because the most associated SNPs are located in intron 19 of the gene (*18-24*). However, we have identified *DEXI* as a causal candidate gene in the 16p13.13 region our previous work, using expression quantitative trait locus (eQTL) mapping and chromosome confirmation capture (3C) in human monocytes. The DEXI protein is predicted to be 10kDa in size, with a potential transmembrane domain but no other clearly defined features. The DEXI gene is highly conserved among species, including the mouse, and has no paralogue in the human genome.

The *DEXI* promoter is 150 kb away from the T1D-associated SNPs (*25*). We have proposed that the disease-associated SNPs affect disease risk by altering transcription factor binding to the *DEXI* promoter, which we have shown to be physically linked to intron 19 of *CLEC16A* by a chromatin loop. We have also demonstrated that increased T1D risk mediated through 16p13.13 is associated with decreased *DEXI* expression, implying that wild-type or normal *DEXI* function has a protective function in autoimmune disease (*25*). Our eQTL result has since been replicated by other groups (*24, 26*), but in the absence of a *Dexi* knockout mouse model, definitive studies to extend evidence in support of *DEXI*’s preventive role in T1D have not been possible. Furthermore, there are so far no known rare variants of *DEXI* in humans with informative phenotypes.

The NOD mouse develops spontaneous T1D from around 12 weeks of age, with approximately 75% of females affected by 30 weeks of age (*27*). The disease is characterized by lymphocytic pancreatic islet infiltration (insulitis) and progressive autoimmune destruction of the pancreatic beta cells, which is evident from 6-8 weeks of age (*27*). Here, in agreement with our prediction that *DEXI* has a protective role in T1D, we report that disruption of *Dexi* in the NOD mouse leads to accelerated spontaneous autoimmune diabetes when compared to the wild-type NOD. Furthermore, we show that *Dexi* disruption alters the host metabolome, gut microbiome, and serum IgA and IgM levels in a manner consistent with a role for *DEXI* in influencing the interaction between the immune system and microbiota. Our results support the role of *DEXI* as a causal factor in human T1D risk, related to a direct or indirect effect on the production of microbial metabolites.

### *Dexi* disruption in NOD mice does not alter breeding fitness or histological phenotype

We used CRISPR-Cas9 genome-editing technology in NOD zygotes to introduce disruptive mutations in the coding sequence of *Dexi* in NOD offspring. Two mutations contained entirely within the *Dexi* protein-coding sequence were selected for study. These mutations were a one base-pair insertion (*INS1BP*) and a twelve base-pair deletion (*DEL12BP)* (**Figure 1A and 1B; Supplementary Figures S2A and S2B; Supplementary Table S2**), occurring early in the protein coding sequence. The *INS1BP* mutation has the potential to cause a more severe phenotype than the *DEL12BP* mutation since it causes a frame shift and nonsense transcript, whereas the 12-bp deletion removes 4 amino acids but leaves the rest of the protein in frame.

**Figure 1.**
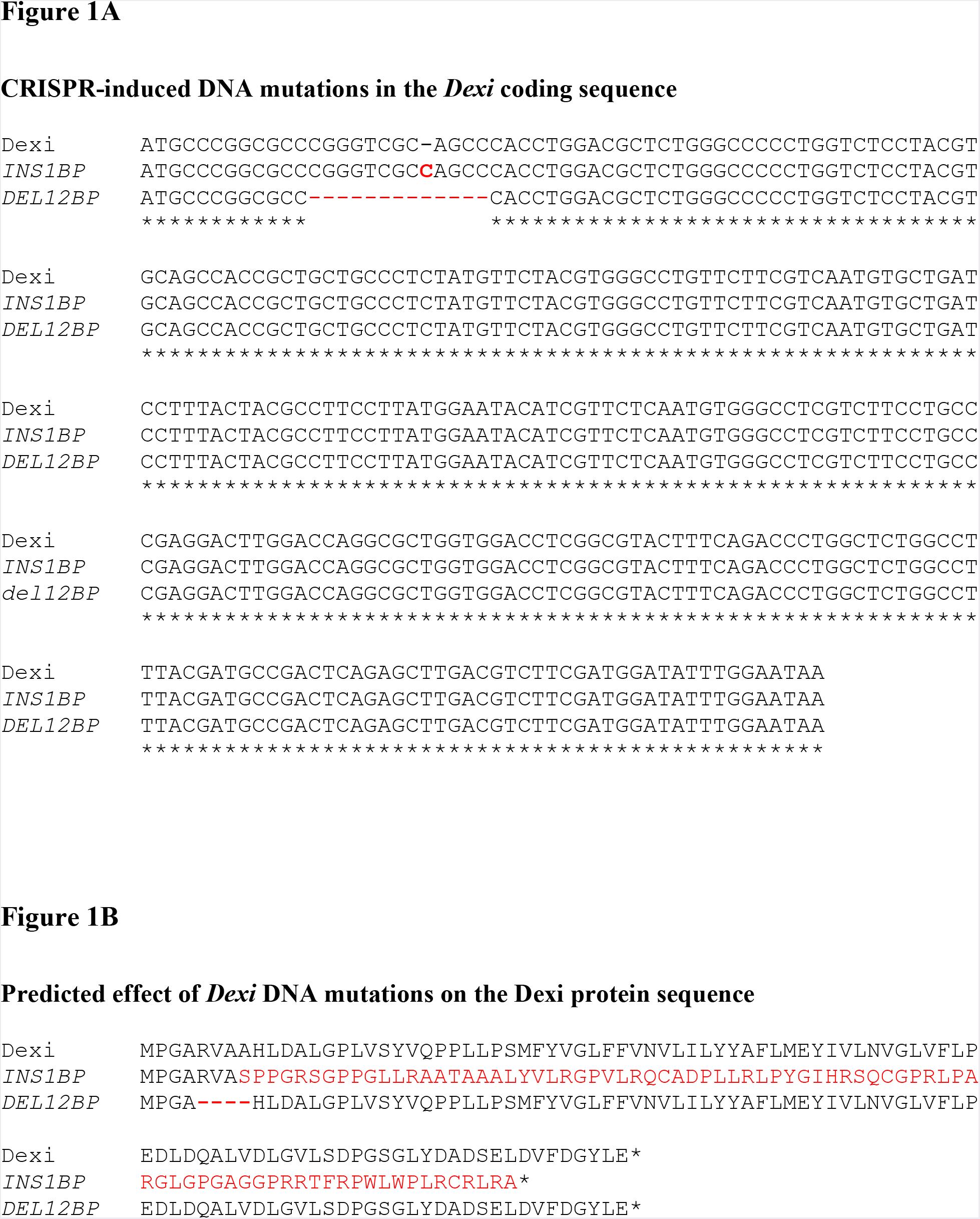
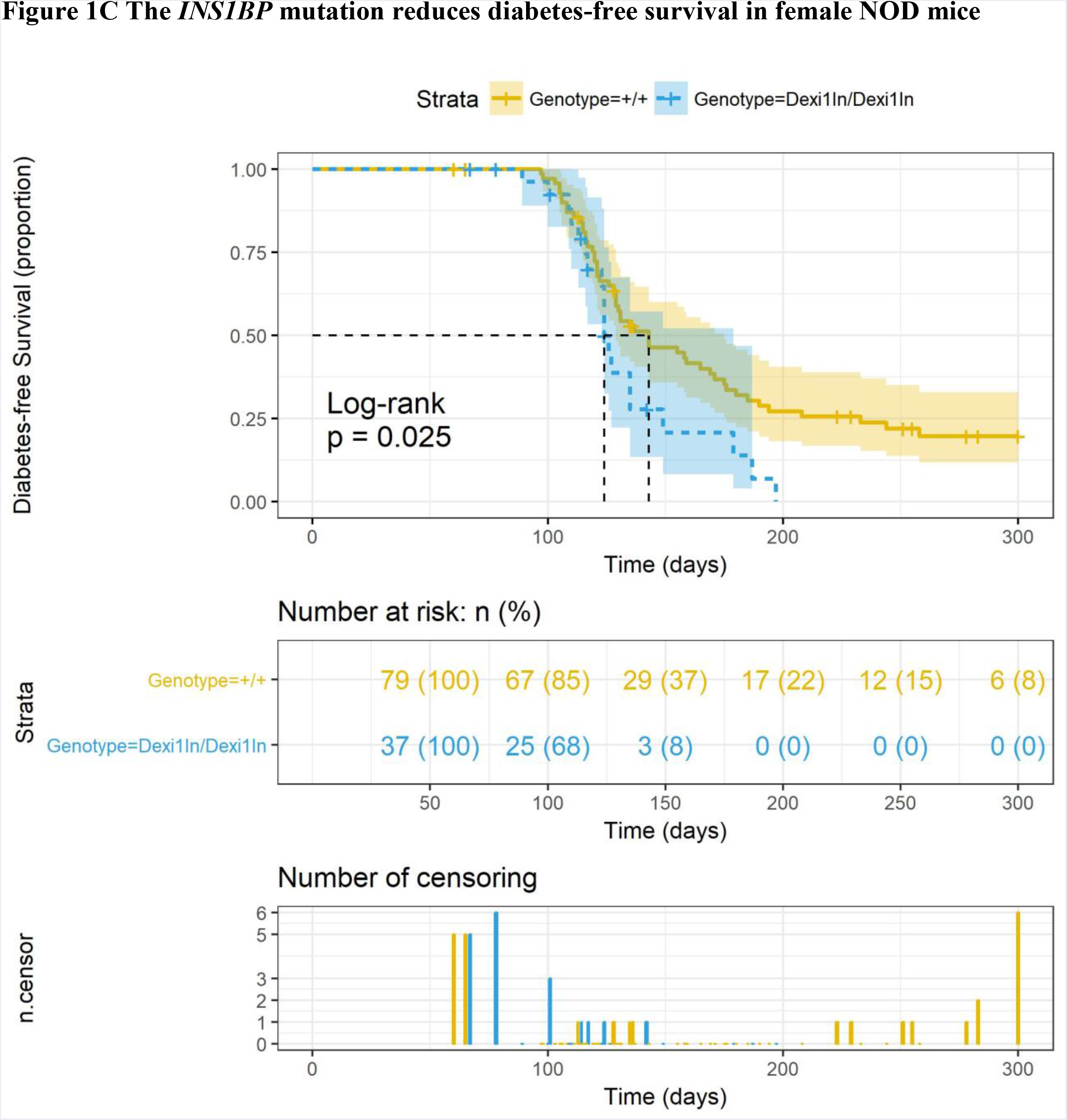
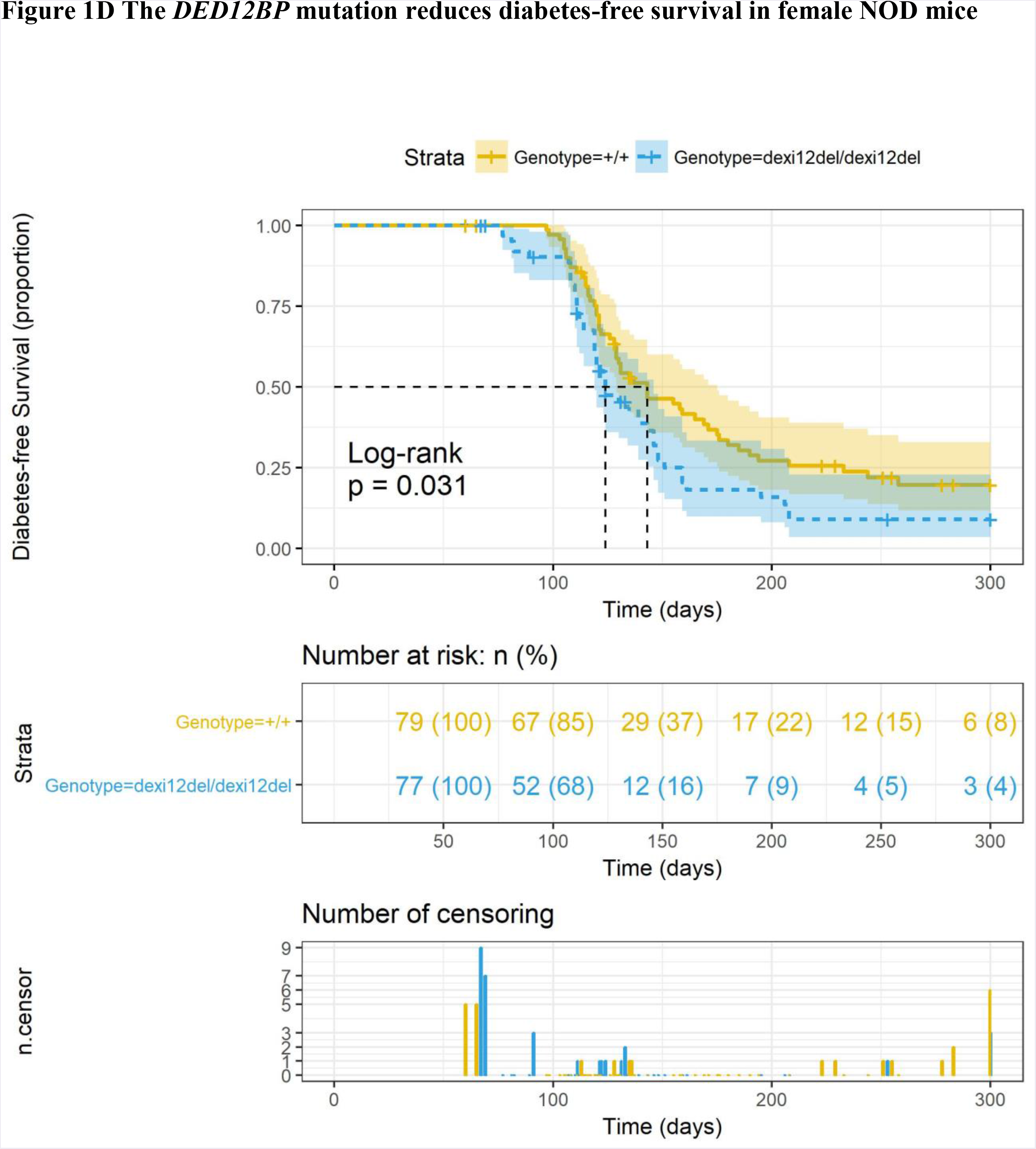
CRISPR induced mutations in Dexi gene and predicted effect on protein sequence. **A: The *INS1BP* and *DEL12BP* CRISPR-induced mutations in the *Dexi* DNA coding sequence**. Mutations are shown in red. **B: The predicted impact of the *INS1BP* and *DEX12BP* mutations on the Dexi protein**. Changes in the amino acid sequence are shown in red. **C and D: The presence of a homozygous *INS1BP* (C) or *DEL12BP* (D) *Dexi* mutation reduces diabetes-free survival in female NOD mice. Top panel for each line:** Kaplan-Meier diabetes-free survival curve with log-rank p-value for wild-type NOD mice (n=79) represented as yellow-orange (+/+). *Dexi*-disrupted mice survival curves are represented in blue, for homozygous *INS1BP* mice (n = 37; labelled as Dexi1in/Dexi1in) and *DEL12BP* mice (n=77); labelled as dexi12del/dexi12del). 95% Confidence Intervals are shown for each genotype as translucent shaded regions surrounding the genotype line. **Mid panel:** Numbers of animals at risk over time. **Bottom panel:** Bar plot of censorship data for each line.

Homozygous mice were viable and the mutations had no apparent impact on breeding fitness. Littermate wild-type NOD and homozygous NOD mice carrying the *INS1BP* or *DEL12BP* mutations were evaluated by histopathology of multiple tissues including pancreas, salivary gland, liver, spleen, kidney, bone marrow, thymus, lung, heart, gastro-intestinal tract and brain. Neither *Dexi* mutation was associated with histologically identifiable changes (representative images are shown in **Supplementary Figure S8**).

In order to establish the effect of the introduced mutation on Dexi protein expression, we first sought to identify the endogenous Dexi protein in wild-type NOD mice using a range of commercially available anti-DEXI antibodies for western blotting of mouse tissue lysates. However, the available commercial antibodies did not allow a conclusive investigation of Dexi protein levels in the mutant NOD mice **Supplementary Figures S2C, S2D and S2E**. Quantitative PCR of thymus and spleen demonstrated that transcripts of the disrupted *Dexi* gene were still detectable in both lines of *Dexi-*disrupted mice, despite the presence of CRISPR-induced mutations (**Supplementary Figure 2F**).

### *Dexi* disruption in NOD mice accelerates diabetes onset in females

A female bias in diabetes incidence in NOD mice has been reported (*27*), in association with hormonally-driven variations in the microbiome (*28, 29*). In agreement with other studies, in our wild-type NOD colony, 75% of females are diabetic (glycosuric) by 30 weeks of age, compared to 50% of males.

In a 300-day diabetes cumulative frequency study, homozygous female mice in both the *INS1BP* and *DEL12BP* lines had accelerated and increased cumulative diabetes incidence compared to wild-type NOD mice without *Dexi* mutations (**Figure 1C and 1D; Supplementary Table S3**), confirming the protective effect of a functional Dexi gene in autoimmune diabetes. There was a trend towards more exaggerated phenotypic changes in the *INS1BP* line, with the potentially more disruptive mutation, compared to the *DEL12BP* line. However, the experiments were not designed to investigate this issue, so numbers were not high enough to demonstrate statistically significant differences between the two *Dexi*-disrupted lines.

In contrast to our findings in females, no acceleration or increase in cumulative diabetes incidence was seen in male mice in the *INS1BP* and *DEL12BP* lines (**Supplementary Figures S3A and S3B**). This result in males could reflect reduced power due to the lower incidence of diabetes in male wild-type NOD mice compared to females, however no trend towards a difference was observed to suggest this is the case.

### *Dexi* disruption in the NOD mouse does not influence the presence of insulitis

To investigate whether the acceleration and increase in diabetes incidence in *Dexi-*disrupted female mice was associated with increased lymphocytic infiltrate in the islets, blinded histopathology of the pancreas was undertaken in 6-week old mice (**Supplementary Table S4; Supplementary Figure S4**). Multiple tissue sections and islets were cut and examined from each mouse and no significant difference in insulitis could be identified between wild-type NOD and *Dexi*-disrupted mice. This suggests that if *Dexi* disruption affects infiltrating lymphocytes, it alters their function or kinetics of the infiltration rather than their steady state presence or absence in the islets.

Insulin autoantibodies (IAA) were measured as a marker of autoimmunity in serum from in age-matched female *Dexi-*disrupted NOD and wild-type NOD mice (**Supplementary Table S6**). Two of 12 wild-type NOD mice were positive compared to 4 of 12 in each of the Dexi-disrupted lines (**Supplementary Figure S7**). Whilst not statistically significant, this trend is consistent with the finding of accelerated diabetes in NOD mice following disruption of the Dexi gene.

### *Dexi* disruption in NOD mice affects serum IgA and IgM but not cellular immune-phenotype

Since T1D is an immune-mediated condition, to assess the impact of *Dexi* disruption on the cellular immune system, flow cytometric analysis of thymus, spleen and bone marrow in wild-type and *Dexi*-disrupted mice was undertaken using a panel of B-cell, T-cell and myeloid lineage markers (**Supplementary Table S5**). After accounting for multiple tests and hypotheses, no consistent significant differences were detected in cellular immune-phenotype between *Dexi-*disrupted and wild-type NOD mice (**Supplementary Figures S5A to S5F**). To assess the impact of *Dexi* disruption on the humoral immune system, serum IgG, IgM and IgA were evaluated in age-matched female *Dexi-*disrupted NOD and wild-type NOD mice (**Supplementary Table S6**), prior to the onset of the diabetic phenotype. A significant increase in serum IgA (p = 0.0102) and IgM (p = 0.0393) was evident in *Dexi*-disrupted compared to wild-type NOD mice (**Figure 2**).

**Figure 2:**
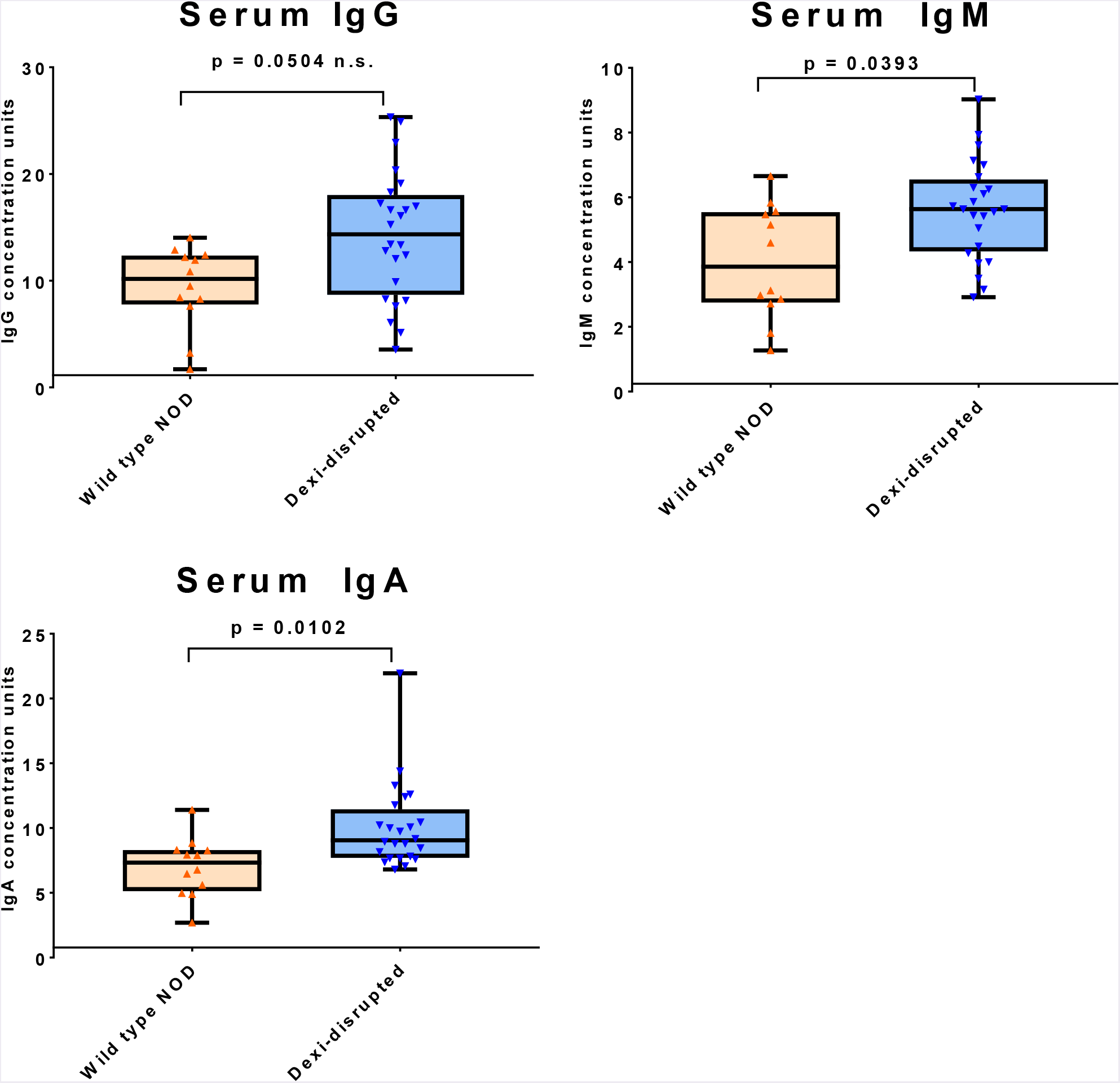
Dexi-disrupted NOD mice demonstrate increased serum IgA and IgM compared to wild-type NOD mice. Serum IgG, IgM and IgA were measured by ELISA in age-matched nondiabetic female *Dexi*-disrupted (blue; n=24) and wild-type (orange; n=12) NOD mice by ELISA and quantified against a standard curve. Groups were compared using the Mann-Whitney test and p values are shown after Bonferroni correction for multiple testing.

The most striking finding was the elevation in serum IgA identified in *Dexi-*disrupted mice. Immunoglobulin A is secreted in the gut, and has multiple functions in mucosal immunity (*30*). Both circulating IgA deficiency and human combined immunodeficiency are associated with variants in the 16p13.13 region of the genome (*15, 16, 31*), so *DEXI* may play a role in IgA biology. Due to the high degree of linkage disequilibrium in the 16p13.13 region, multiple non-coding variants have been associated with both T1D and IgA deficiency. As the direction of risk for T1D and IgA deficiency is the same at 16p13.13 (*15*), the raised serum IgA in association with diabetes risk in this study is surprising.

The regulation of IgA production is complex and multifactorial, and as well as endogenous host factors, IgA production relates to the presence of microbiota in the intestine and other mucosal surfaces (*32-34*). The relationship between IgA and the microbiome is likely to be bi-directional, since microbiota influence IgA production, but the presence and repertoire of mucosal IgA can influence which organisms are able to thrive in the gastrointestinal tract (*35*). D*exi* dosage may also be important here, since the *Dexi* disruption afforded by CRISPR in our model is much more severe than the effect typically seen with a common human allele.

### The fecal microbiome of NOD mice is affected by Dexi-disruption

Gut microbiota influence the immune system and development of diabetes in the NOD mouse (*28, 29, 36-39*), as well as susceptibility to T1D in young children (*40-43*). For example, in the NOD model, mice lacking Toll-like Receptor (TLR) signaling, due to absence of the MyD88 adapter protein, are protected from diabetes in the presence of gut microbiota, but protection is not observed in germ-free mice (*44*). Reconstitution of the gut flora of these germ-free mice with specific bacterial lineages restores the diabetes protection afforded by MyD88 deficiency (*45*). Furthermore, female NOD mice can be protected from the development of diabetes by gavage transfer of male NOD mouse fecal microbiota (*29*). Our finding of diabetes acceleration in *Dexi*-disrupted female but not male NOD mice would be consistent with a sex-influenced microbiome-dependent protective effect of the Dexi gene (*28, 29, 46*).

It has previously been shown that the NOD genetic background is associated with distinct fecal microbiota compared to B6 mice (*47*). Recent evidence also suggests that a small proportion of microbial diversity in the human gut is driven by the host genome (*48*). Since the 16p13.13 region has previously been associated with microbial diversity in the human gut (*17*), and diabetes-associated alleles in other genomic regions have been shown to influence the microbiota in the human gut (*47*), we investigated whether *Dexi*-disruption was associated with changes in the gut microbiome in the NOD mouse.

We examined the fecal microbiome in age-matched (4 weeks, 6 weeks and 10 weeks) female mice across multiple litters and cages (**Supplementary Table S7**), prior to the onset of diabetes. Normoglycemic mice were used for this experiment to avoid any potential consequential impact of the accelerated diabetes phenotype in the *Dexi*-disrupted NOD mouse on the fecal microbiome. DNA was extracted from fecal samples from 44 mice and applied to a microarray designed to detect all known microorganisms in a sample, with species- and strain-level identification **(Supplementary Tables S8, S9, S10).** Eight microbial species showed significantly altered representation in homozygous *Dexi-*disrupted mice compared to wild-type NOD mice (**Figure 3 and Supplementary Figures S7A and S7B**).

**Figure 3:**
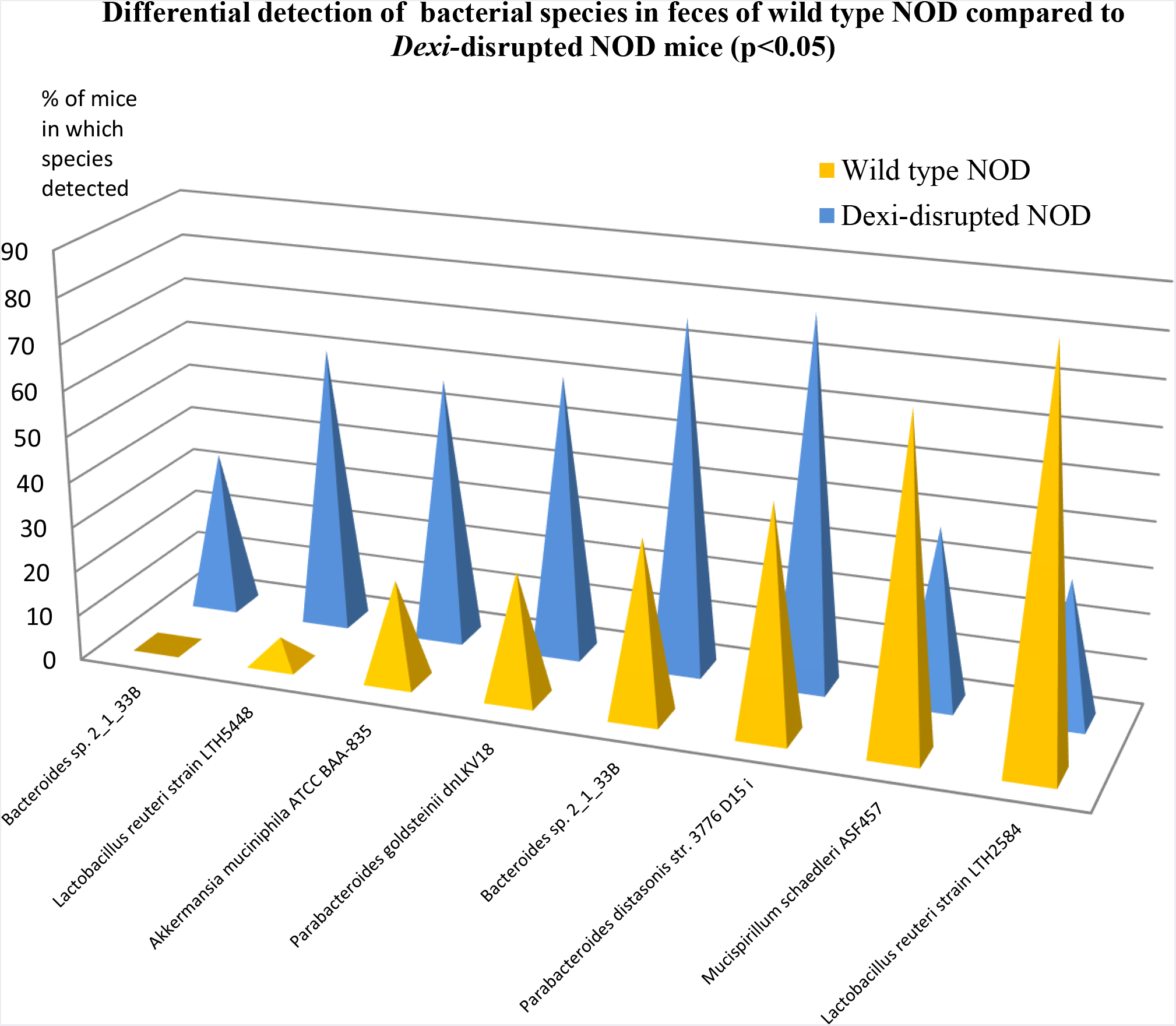
The fecal microbiome of *Dexi-*disrupted NOD mice is different to that of wild-type NOD mice. Bar plot showing robustly detected microbial species on a microbiome array with a significantly different prevalence in DNA extracted from the feces of *Dexi*-disrupted mice compared to wild-type NOD mice. Blue peaks correspond to *Dexi*-disrupted mice (n=26), orange peaks correspond to wild-type NOD mice (n=18).

Several of the microbial species that were present in a significantly different proportion of *Dexi-*disrupted NOD mice compared to wild-type NOD mice have been implicated in human T1D risk. For example, we found significantly higher representation of the *Bacteriodetes* phylum in *Dexi*-disrupted NOD mice, including *Bacteroides*, *Parabacteroides distasonis*, and *Parabacteroides goldsteinii*. Notably, children with T1D have been shown to have a significant increase in the number of *Bacteroidetes* as well as *Bacteroides* (*49*). We also found inverse representation between two strains of *Lactobacillus reuteri*, with *Dexi-*disrupted mice being low in strain LTH2584 (31% vs WT 89%) and high in strain LTH5448 (62% vs WT 6%). *L. reuteri* is an established and widely used probiotic species in research and clinical trials (*50*). It is also notable that *Akkermansia muciniphilia,* a mucus-degrading bacteria detected predominantly *Dexi-* disrupted mice, is one of the microbial species consistently targeted by mucosal IgA in the gut (*33*).

The mechanism(s) by which microbiota impact on diabetes risk are unclear, but may relate to the production of bacterial metabolites which influence the function of immune cells, such as regulatory T cells (*51*).

### *Dexi* disruption in NOD mice leads to metabolic changes in liver, spleen and blood

To investigate whether the metabolomic profiles of *Dexi-*disrupted and wild-type NOD mice differ, particularly with respect to microbial metabolites, we undertook extensive metabolomic and lipidomic analyses of 6-week-old female mice. Mice of this age were chosen to ensure that any changes were related to *Dexi* disruption rather than an early diabetic phenotype. In addition to whole blood, we examined spleen because of its central role in the immune system and liver because it receives a blood supply directly from the gut, via the hepatic portal vein. The levels of multiple non-lipid low molecular weight metabolites differed between *Dexi-*deficient and wild-type NOD mice (**Figure 4A**). Liver showed the highest number of significant changes, but a proportion of these changes were also seen in spleen and blood. These differences were not associated with changes in the histological appearance of the liver in any mice (**Supplementary Figure S8**). There was no significant difference in lipid profile among the groups of mice in blood, spleen or liver.

**Figure 4:**
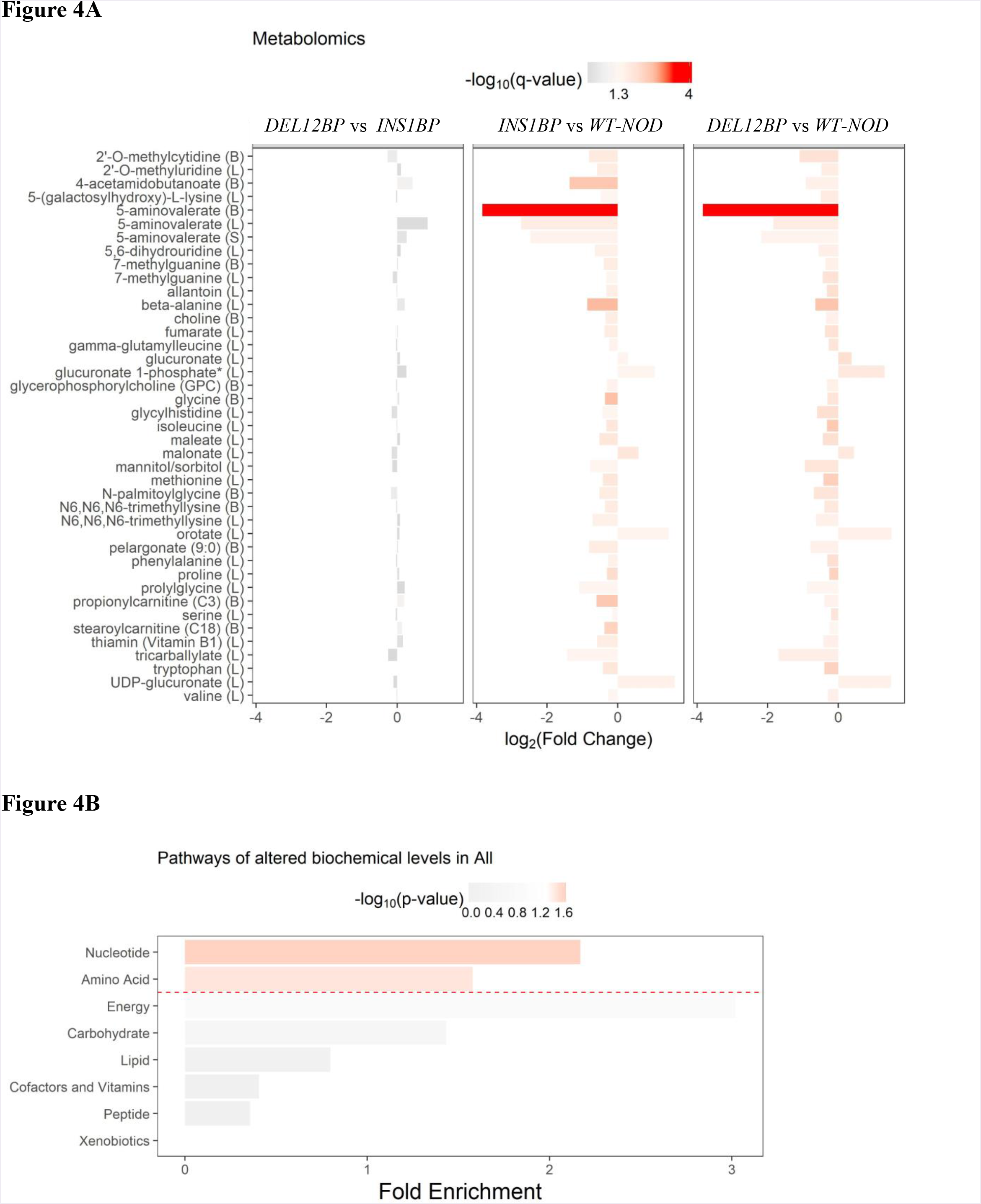
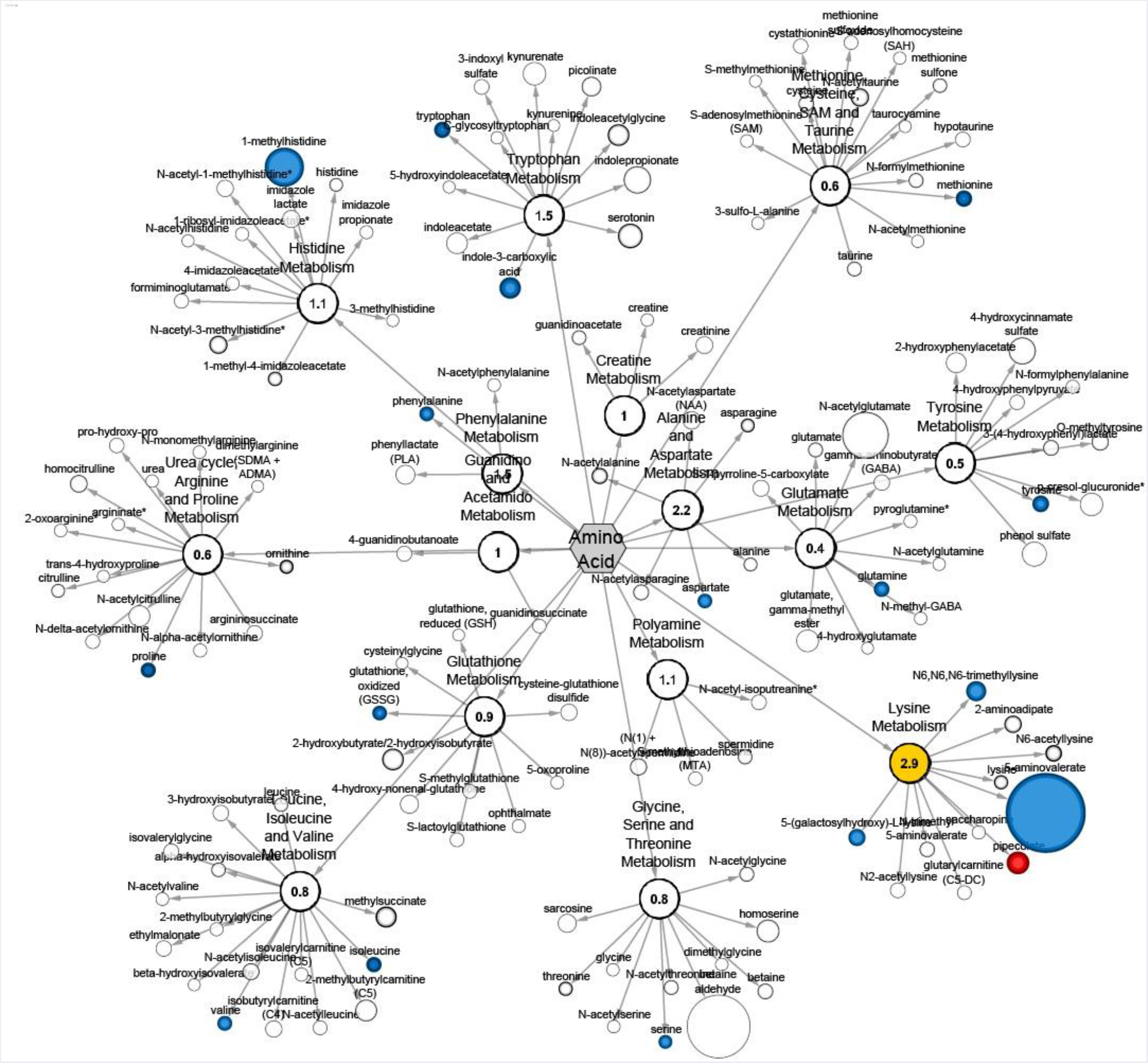
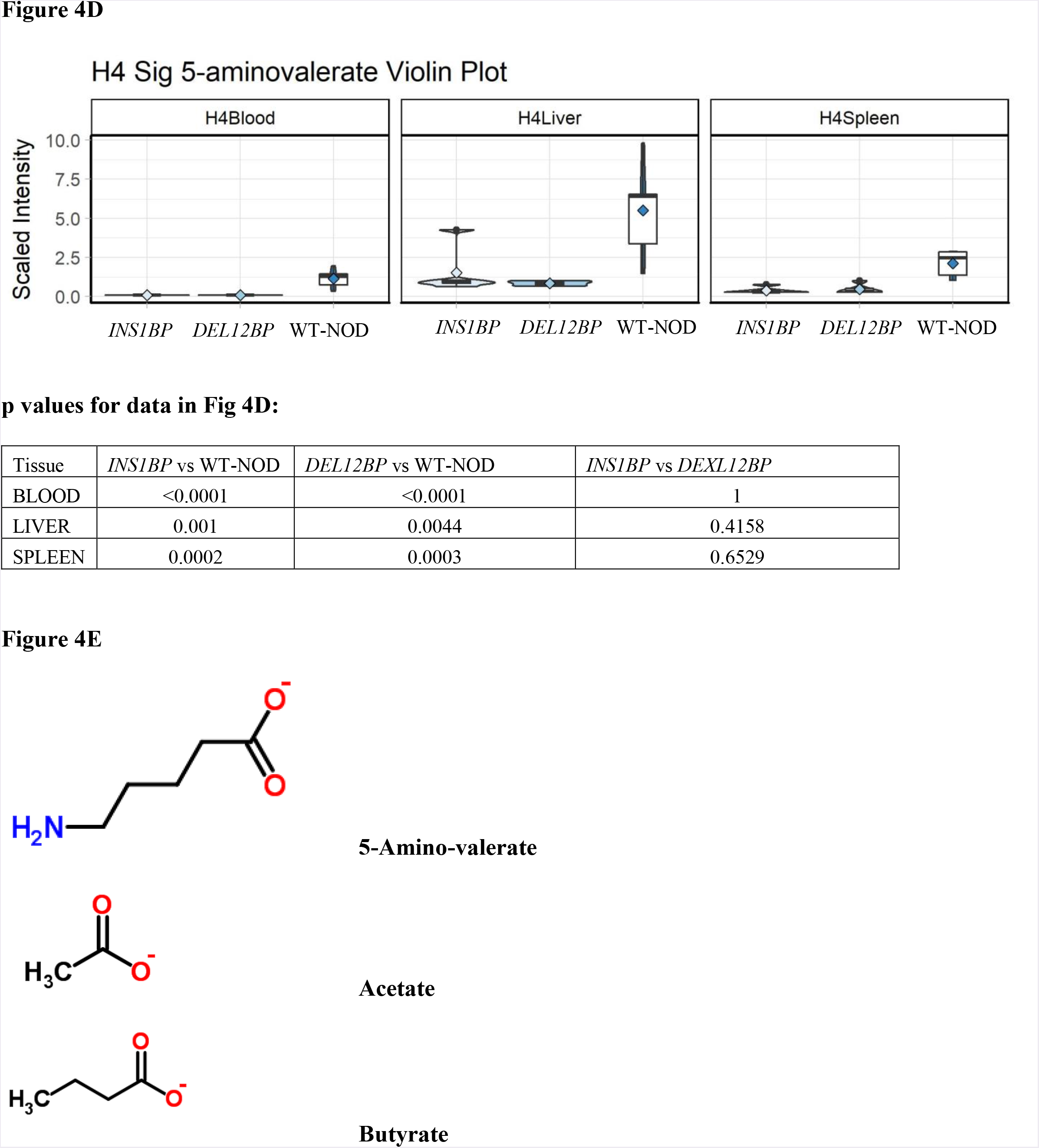
Metabolomic changes in *Dexi*-disrupted NOD mice compared to wild-type NOD mice. **A. Bar plot of biochemical compounds demonstrating significant and consistent fold-change direction in 6-week old females from both *INS1BP* and *DEL12BP Dexi*-disrupted mouse lines (n=5 per line) compared to wild-type NOD (WT) (n=5).** Bar color represents -log10(q-value) scale, where red is significant and grey is not significant. The x-axis is log2(Fold Change) of the biochemical level in *Dexi*-disrupted compared to wild-type NOD mice. Parentheses beside the biochemical name on the y-axis indicate sample tissue type: (L) = Liver, (B) = Blood and (S) = spleen. **B. Pathway set enrichment analysis of compounds in A.** The red horizontal dashed line separates non-significant pathways (grey) from significant pathways (red), where bar color is scaled by -log10(p-value). **C. Amino acid pathway network (Cytoscape) for mouse liver comparing *INS1BP* to wild-type NOD mice.** Shaded nodes correspond to significantly different metabolites (blue for reduced in *Dexi*-disrupted *INS1BP* mice; red for increased in *Dexi*-disrupted mice; yellow for a significant change in a pathway). Dark red or blue represent change at p < 0.05 and lighter colored nodes correspond to trending threshold p < 0.1. White nodes correspond to detected compounds with no significant change. **D. Biochemical scaled intensity violin plots of 5-aminovalerate in liver, blood, and spleen in *Dexi-*disrupted lines and WT-NOD mice shown in A. E. Schematic structures of 5-aminovalerate and its breakdown products acetate and butyrate**

The metabolomic analysis demonstrated that the two pathways most significantly and consistently impacted by *Dexi* disruption were those of nucleotide synthesis and amino acid metabolism (**Figure 4B**). Several compounds associated with pyrimidine and purine metabolism were consistently depleted in 6-week-old *INS1BP* and *DEL12BP* mice, including allantoin, methyluridine, dihydrouridine, beta-alanine and 7-methylguanidine, with only one, oronate, being consistently elevated. In addition, several amino acids and their associated metabolites were significantly lower in the liver (**Supplementary Figure S9A**) and blood (**Supplementary Figure S9B**) of both *INS1BP* and *DEL12B* mice, consistent with *Dexi*-associated changes in protein metabolism. At the pathway level, several metabolites within the lysine metabolism pathway were affected in both *INS1BP* and *DEL12BP* mice (**Figure 4C; Supplementary Figures S10A to S10C**). One such compound, 5-amino-valerate, was reduced in liver, spleen and blood in both *INS1BP* and *DEL12BP* mice compared to wild-type NOD (**Figure 4D**). This branched chain fatty acid (BCFA) compound is a product of protein degradation by anaerobic bacteria, particularly *Clostridial* strains, and is known to be absent from the metabolome of germ-free mice (*52, 53*). We detected this compound at the highest concentration in the liver, which receives the ‘first-pass’ blood from the gastro-intestinal tract via the hepatic portal vein, providing further evidence for this metabolite being produced by bacteria in the gut.

BCFAs can influence immunity by modulating the host mucosal immune system and impacting on signaling pathways in epithelial cells (*54, 55*). BCFAs can also affect bacterial gene expression of enzymes associated with amino acid metabolism (*56*). Notably, further degradation of 5-amino valerate produces ammonia and the other volatile short chain fatty acids (SCFAs) acetate and proprionate **(Figure 4E)** (*53*). These SCFAs which can be derived from 5-aminovalerate protect against type 1 diabetes in NOD mice by decreasing the frequency of autoreactive T cells in lymphoid tissues, through effects on B cells and regulatory T cells (*57*).

To determine whether depletion of SCFA precursors in the blood and liver is associated with depletion of acetate, butyrate, isobutryate or proprionate in the *INS1BP* and *DEL12BP* mice, we measured these four compounds in the serum and liver of a second cohort of 6-week old female mice (n=10) (**Figure 5**). Consistent with our hypothesis that SCFA depletion would be associated with *Dexi*-disruption, butyrate was depleted in the liver of the *Dexi*-disrupted NOD mice compared to wild-type NOD mice (p = 0.0016 - **Figure 5a**). There was also a trend for acetate depletion in the *Dexi-*disrupted NOD mice but this was not significant after correction for multiple hypothesis testing. The relative depletion of butyrate in the liver was not evident in serum (**Figure 5b**), which provides further evidence for its production by the microbiota in the gut and subsequent transport via the hepatic portal vein.

**Figure 5:**
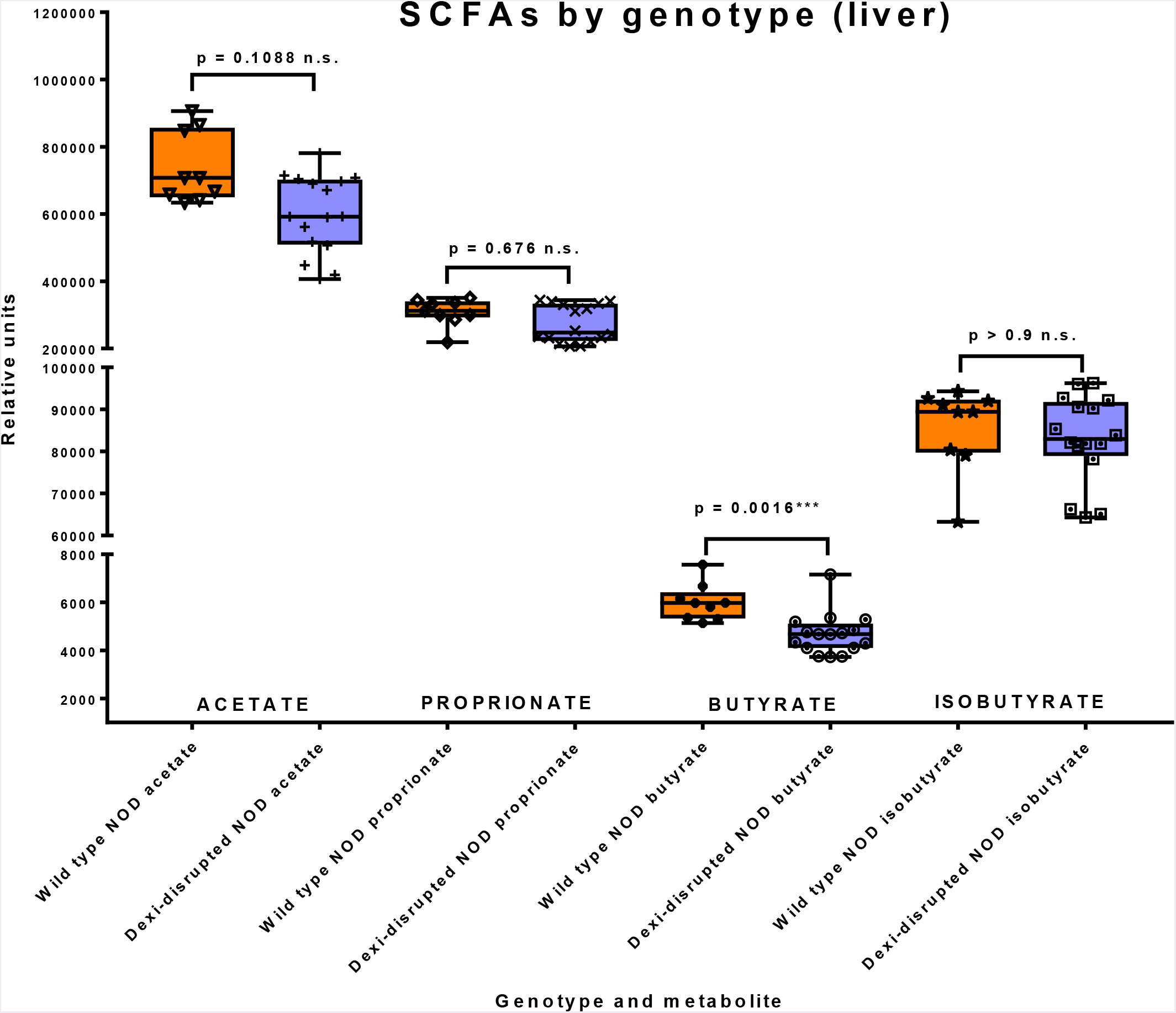
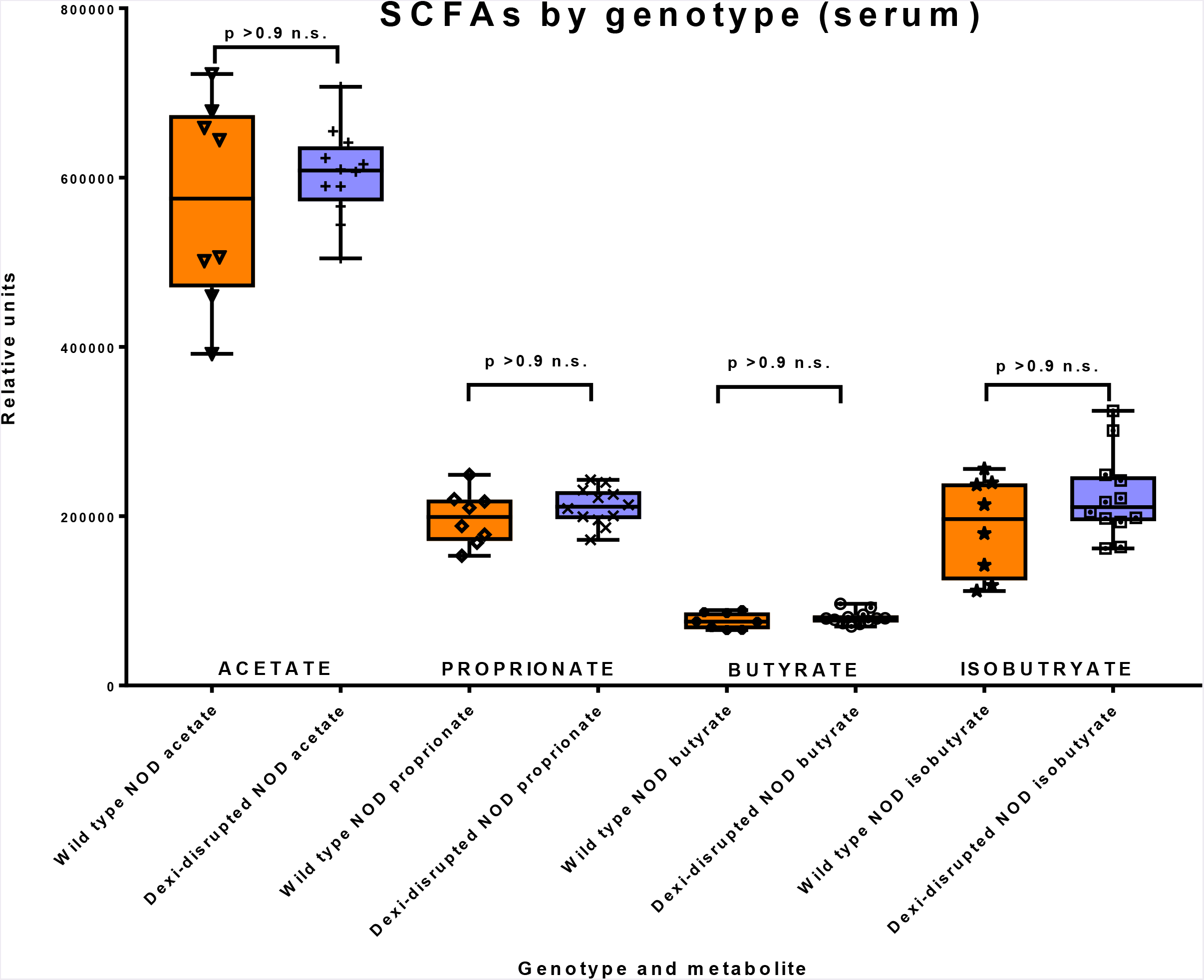
Box and whisker plots demonstrating metabolomic data (short chain fatty acid measurement) from wild-type NOD (n=4) and *Dexi*-disrupted NOD mice (n=6) in liver (a) and serum (b). Groups were compared using the Mann-Whitney test and p values are shown after Bonferroni correction for multiple testing. Technical replicates of each sample were run - (a) triplicate and (b) duplicate; individual data points are shown.

We conclude that *Dexi* has a protective effect in type 1 diabetes. Based on our data, we propose a model whereby *Dexi* disruption alters the gut microbiome, which in turn influences both serum IgA production and diabetes risk via metabolites in pathways including lysine metabolism, BCFAs and SCFAs (**Figure 6**). Our data demonstrate that increased serum IgA (and IgM) are associated with diabetes susceptibility and *Dexi* disruption. We propose that altered microbial metabolism in the gut results in a lack of substrate for synthesis of SCFAs in *Dexi*-disrupted female mice, leading to acceleration of spontaneous diabetes. We hypothesize that the importance of *Dexi’s* effect on the microbiome is reduced in male NOD mice because they naturally harbor a more diabetes-protective microbiome than female NOD mice (*28, 29*). We confirm *Dexi* as a diabetes risk gene which impacts on the gut microbiome and measurable microbial metabolites.

**Figure 6.**
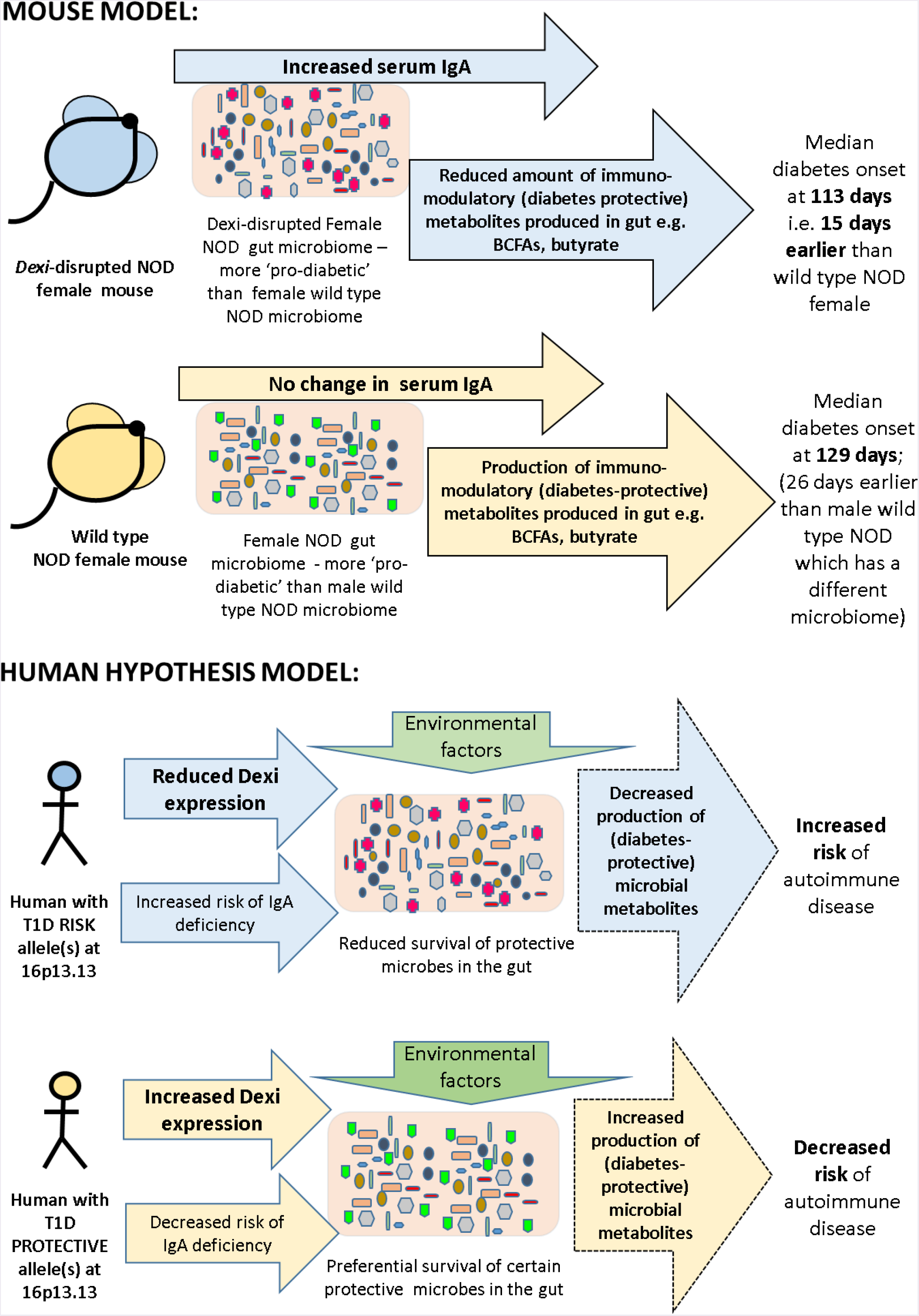
– Summary and proposed model of *Dexi* impact on serum IgA, microbiome, metabolome and diabetes risk

This study has several important implications. Firstly, it illustrates the importance of tissue-specific metabolic studies for identification of disease-associated metabolic phenotypes, rather than reliance on serum measurements. Secondly, it demonstrates an approach by which a candidate causal gene of unknown function, identified by a GWAS study, may be confirmed in the context of a specific complex disease. Thirdly, this study highlights the complex relationship between host genetic variations, serum immunoglobulins (particularly IgA) and microbial diversity in the gut. Finally, our results highlight *DEXI* as a novel therapeutic target in the development of protective strategies in a wide variety of complex autoimmune diseases including T1D, multiple sclerosis and primary biliary sclerosis.

## Supplementary materials

**Supplementary Figure S1:**
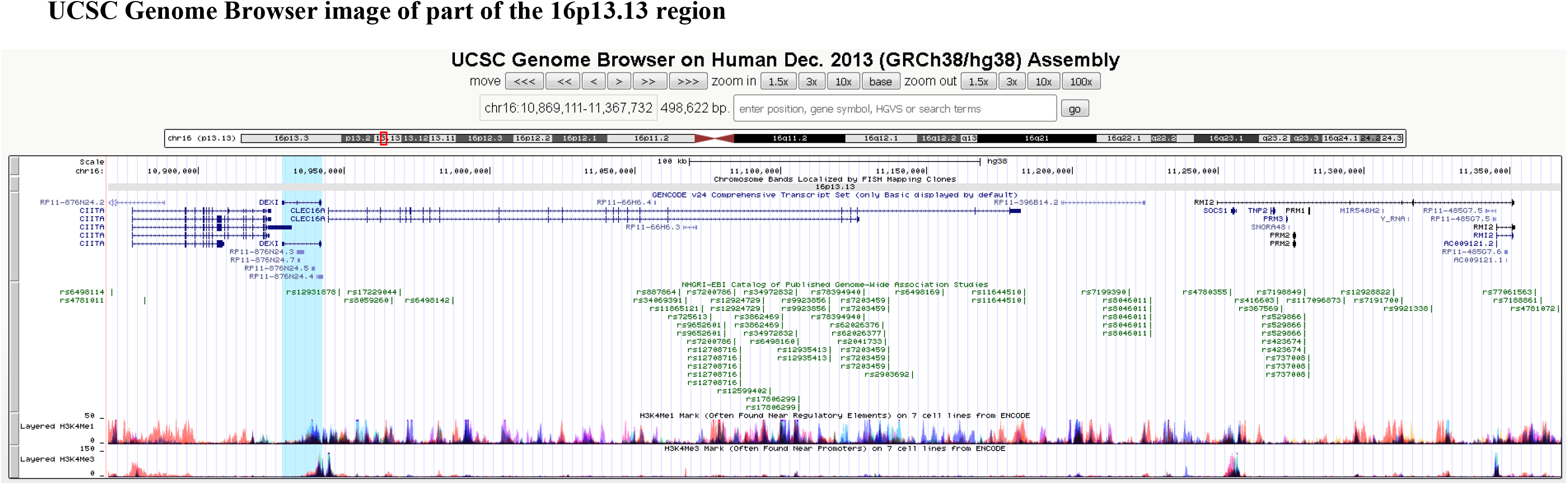
A screenshot from the UCSC genome browser illustrating the relationship between *DEXI* and the autoimmune disease-associated SNPs clustered in and around intron 19 of *CLEC16A*. The DEXI gene is highlighted in blue and published (significant) GWAS SNPs are coloured in green **Supplementary Figure S2A:** Genomic DNA was amplified using gene specific primers and Q5 Hi-fidelity DNA polymerase for 35 cycles. Samples were separated on a 2% agarose gel for 1 hour following a 6-hour digestion with *AvaII* enzyme. Wild-type NOD (WT) do not demonstrate any evidence of digestion, but *INS1BP Dexi*-disrupted mice display either partial digestion (Het – heterozygotes) or full digestion (*INS1BP* Ho – homozygotes) due to the presence of a 1bp insertion which generates an *AvaII* digestion site.

**Supplementary Figure S2A:**
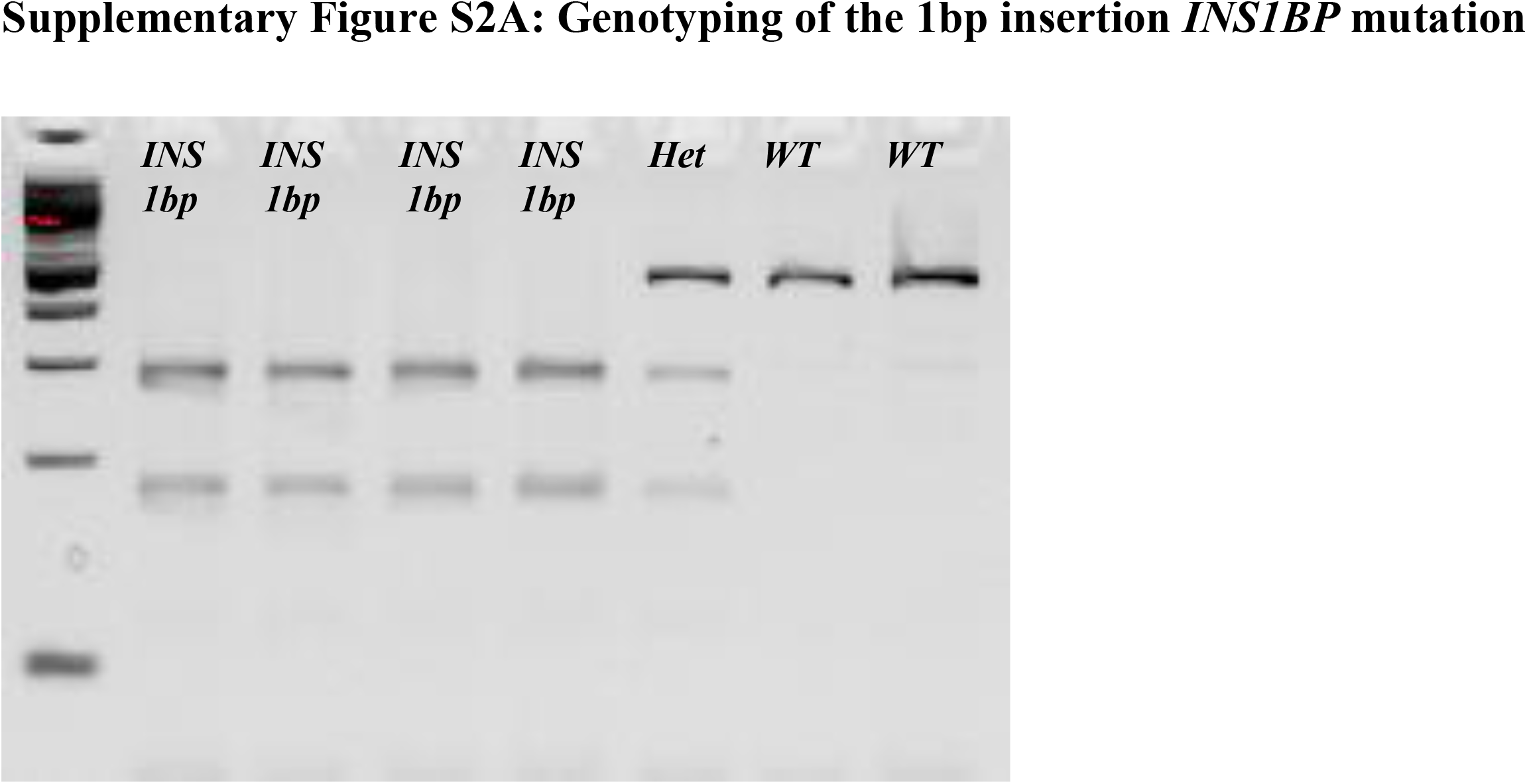
Genomic DNA was amplified using gene specific primers and Q5 Hi-fidelity DNA polymerase for 35 cycles. Samples were separated on a 2% agarose gel for 1 hour following a 6-hour digestion with *AvaII* enzyme. Wild-type NOD (WT) do not demonstrate any evidence of digestion, but *INS1BP Dexi*-disrupted mice display either partial digestion (Het – heterozygotes) or full digestion (*INS1BP* Ho – homozygotes) due to the presence of a 1bp insertion which generates an *AvaII* digestion site.

**Supplementary Figure S2B:**
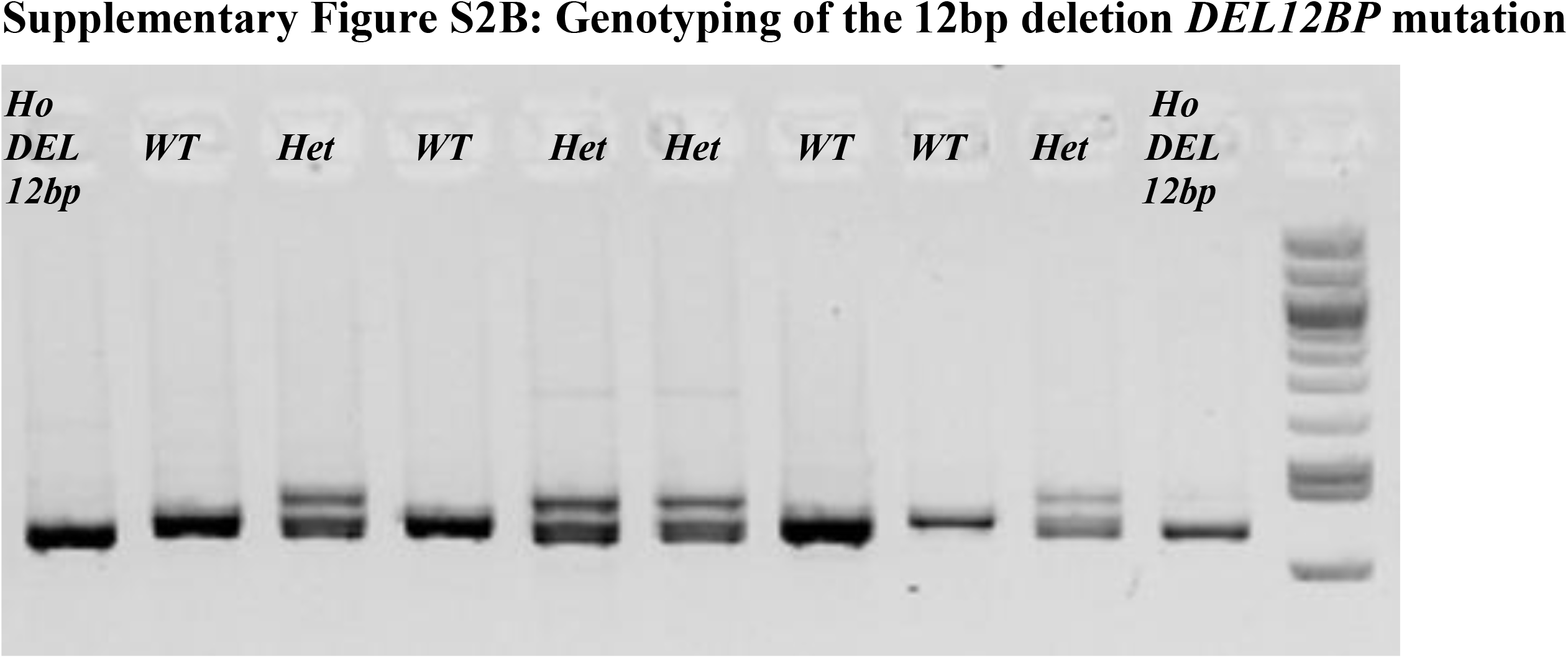
Genomic DNA was amplified using gene specific primers and Q5 Hi-fidelity DNA polymerase for 35 cycles. Samples were separated on a 5% agarose gel for 2 hours at 100V. The amplicon from wild-type (WT) mice is 12bp larger than the amplicon from the *Dexi*-disrupted *DEL12bp* homozygous mice (*Ho*). Heterozygous mice (Het) demonstrate three bands due to heteroduplex formationExperiments to investigate the Dexi protein focused on tissue where Dexi expression has been reported by transcriptomic studies e.g. liver. We identified two antibodies (Sigma Prestige and Proteintech) that were able to recognize recombinant DEXI, giving a single band of the correct size in an over-expression lysate sample. However neither antibody revealed a band corresponding to endogenous Dexi in any mouse tissues and nor were differences between wild-type NOD and Dexi-disrupted mice evident. (Representative western blots are shown in **Supplementary Figures S2C and S2D**). Several other commercial antibodies failed to recognize the recombinant protein and also resulted in multiple non-specific bands when tested against tissue lysate.

**Supplementary Figure S2C:**
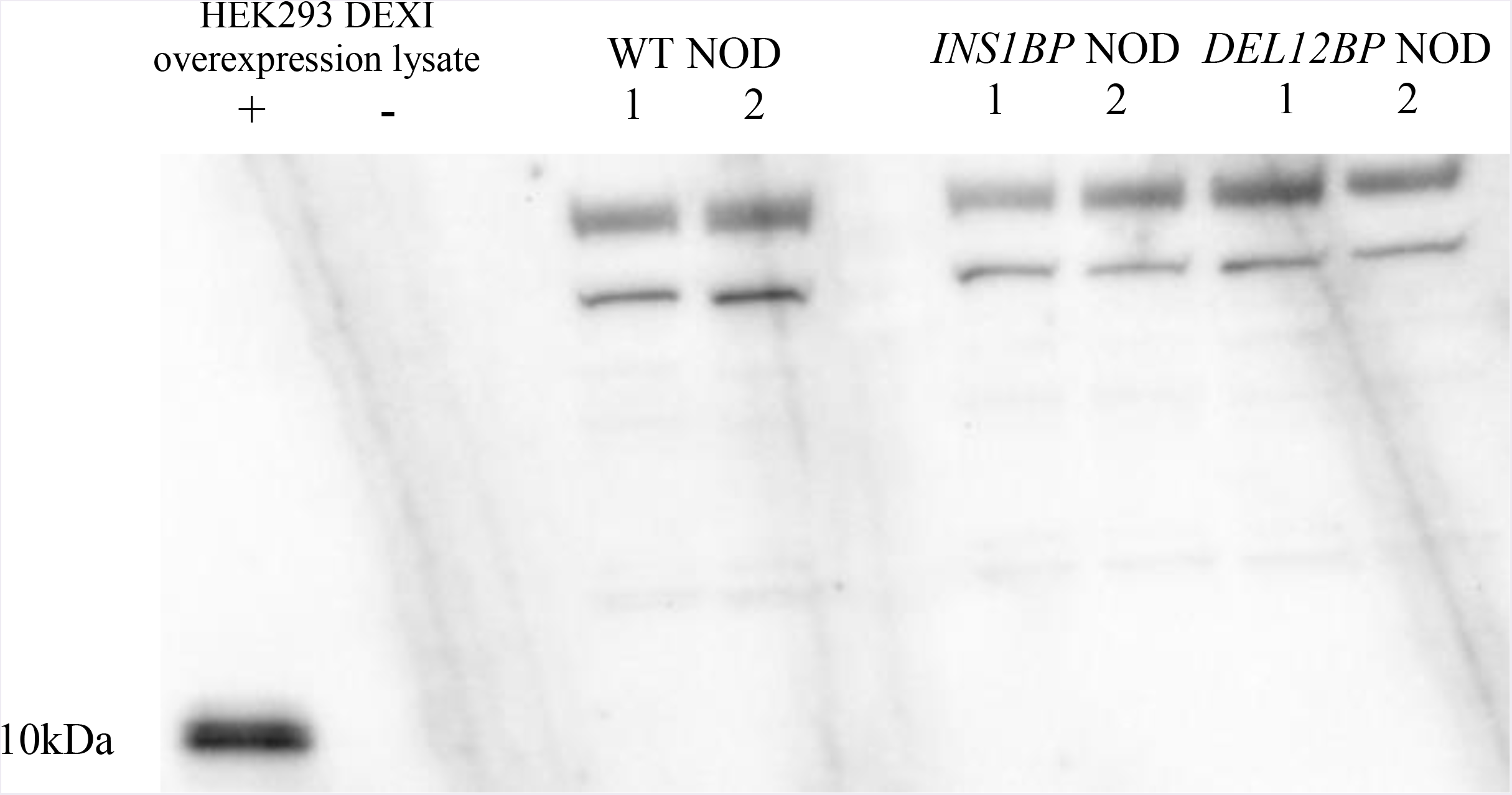
Western blot demonstrating reactivity of commercial polyclonal antibody with recombinant DEXI (+) but no evidence of a band of the correct size in mouse liver lysates (1 and 2) from wild-type NOD (WT NOD) or Dexi-disrupted NOD mice (*INS1BP* and *DEL12BP*). 50ul protein per well was loaded on a 4-20% Tris-Glycine SDS-PAGE gel and blotted onto a PVDF membrane following electrophoresis. The primary antibody for western blotting was a rabbit anti-DEXI polyclonal antibody (Sigma Prestige) at 1:400 dilution, the secondary antibody was a goat anti-rabbit HRP conjugate (VectorLabs) at 1:50,000 dilution and the blot was developed with ECL Advance reagent.

**Supplementary Figure S2D:**
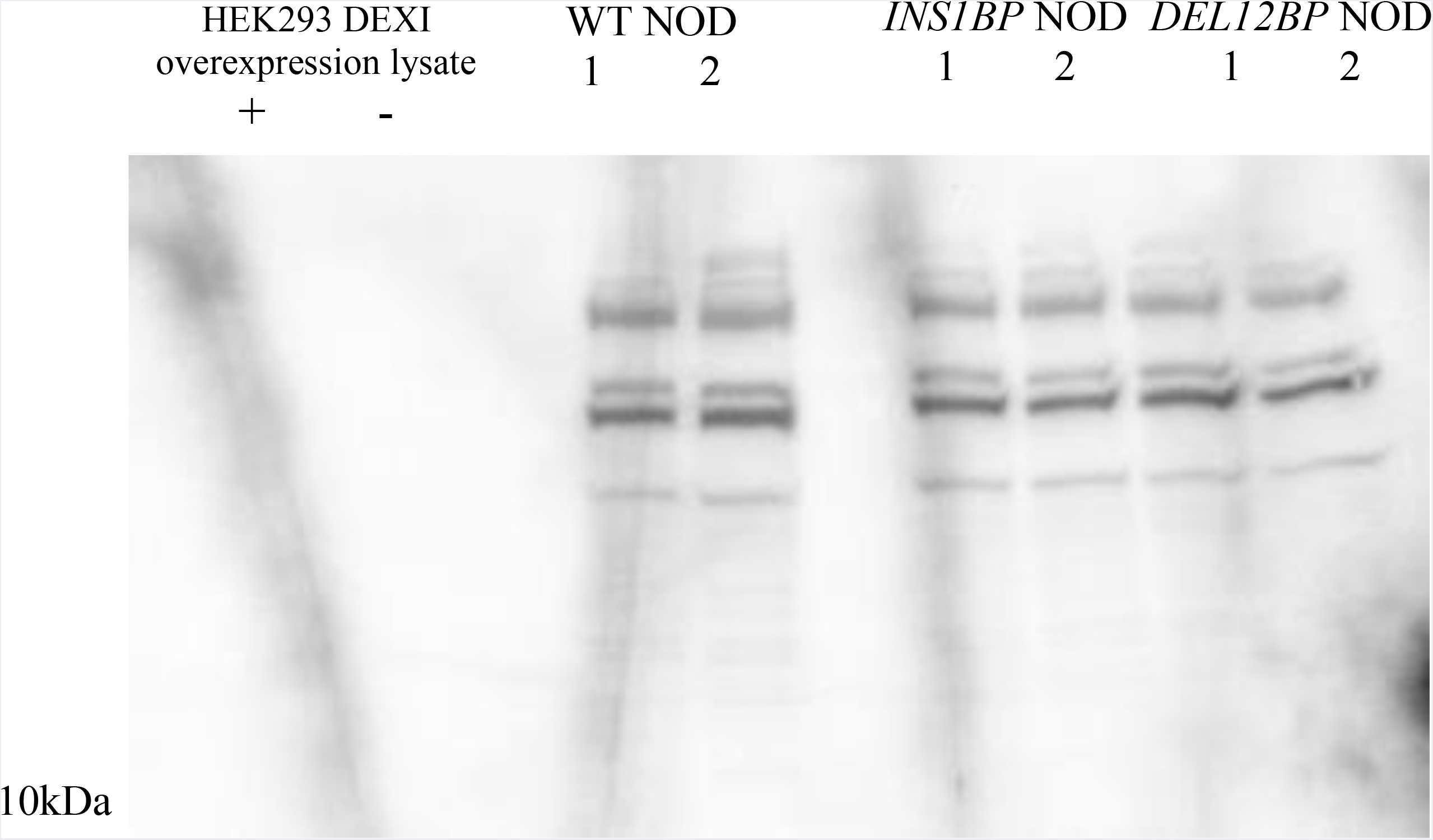
Western blot demonstrating lack of a Dexi band of the correct size using a commercial polyclonal antibody with recombinant DEXI (+) and with mouse liver lysates (1 and 2) from wild-type NOD (WT NOD) or Dexi-disrupted NOD mice (*INS1BP* and *DEL12BP*). 50ul protein per well was loaded on a 4-20% Tris-Glycine SDS-PAGE gel and blotted onto a PVDF membrane following electrophoresis. The primary antibody for western blotting was a rabbit anti-DEXI polyclonal antibody (Abgent) at 1:400 dilution and the blot was developed with ECL Advance reagent.A further attempt to identify Dexi protein using the Sigma Prestige polyclonal antibody was made by immunohistochemistry on wild-type and Dexi-disrupted mouse tissues but a persistent high background staining was observed despite optimization (**Supplementary Figure S2E**).

**Supplementary Figure S2E.**
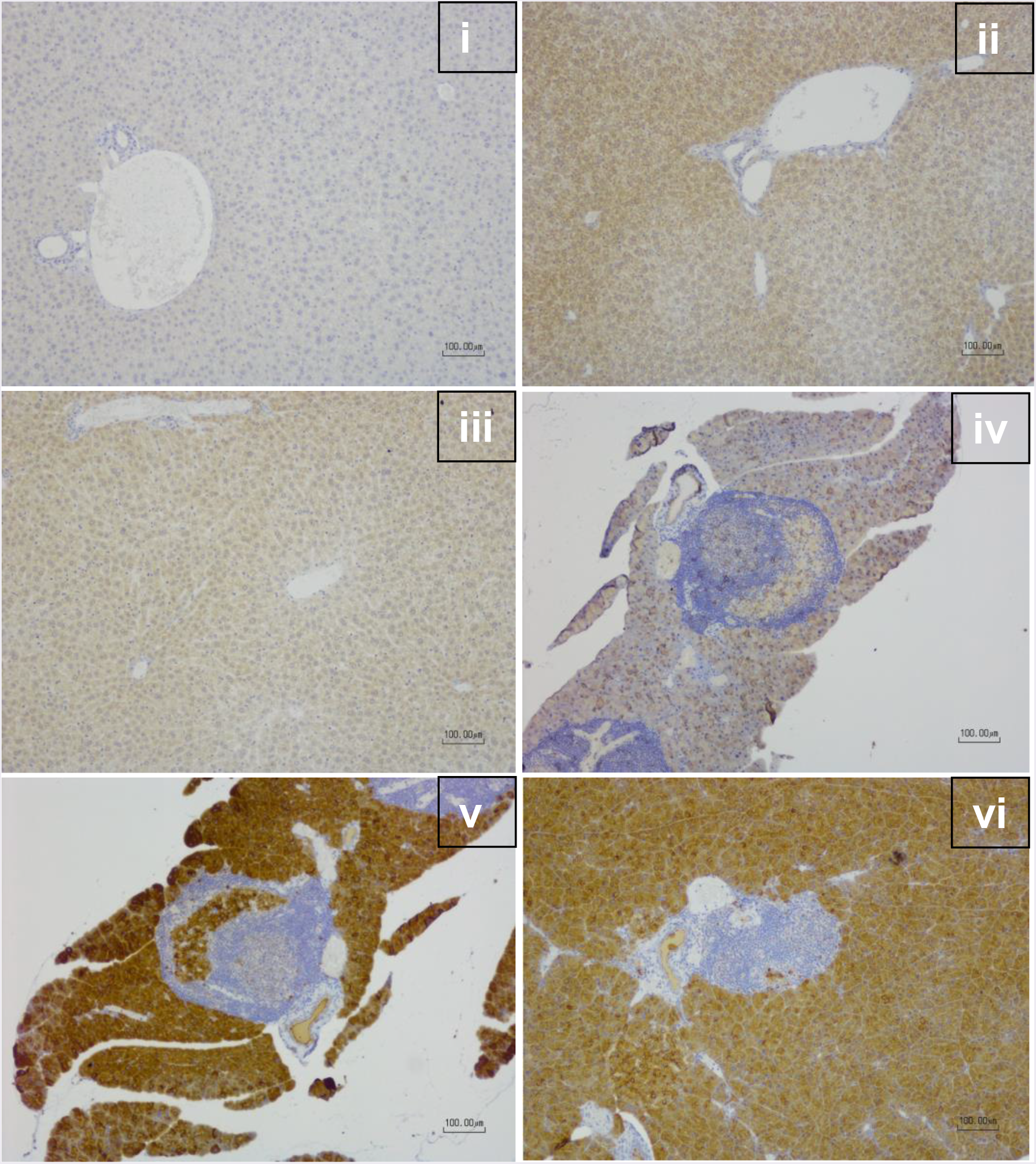
Immunohistochemical staining for DEXI using a commercial polyclonal antibody (Sigma Prestige) reveals non-specific background staining in wild -type NOD mice (ii,v) and *Dexi*-disrupted NOD mice (iii,vi). Representative sections of liver (i,ii,iii) and pancreas (iv,v,vi) are shown. Immunohistochemical staining (ii,iii,v,vi) and isotype-matched immunoglobulin control (i,iv) all with haematoxylin counterstain. Scale bar indicates 100 microns.

**Supplementary Figure S2F.**
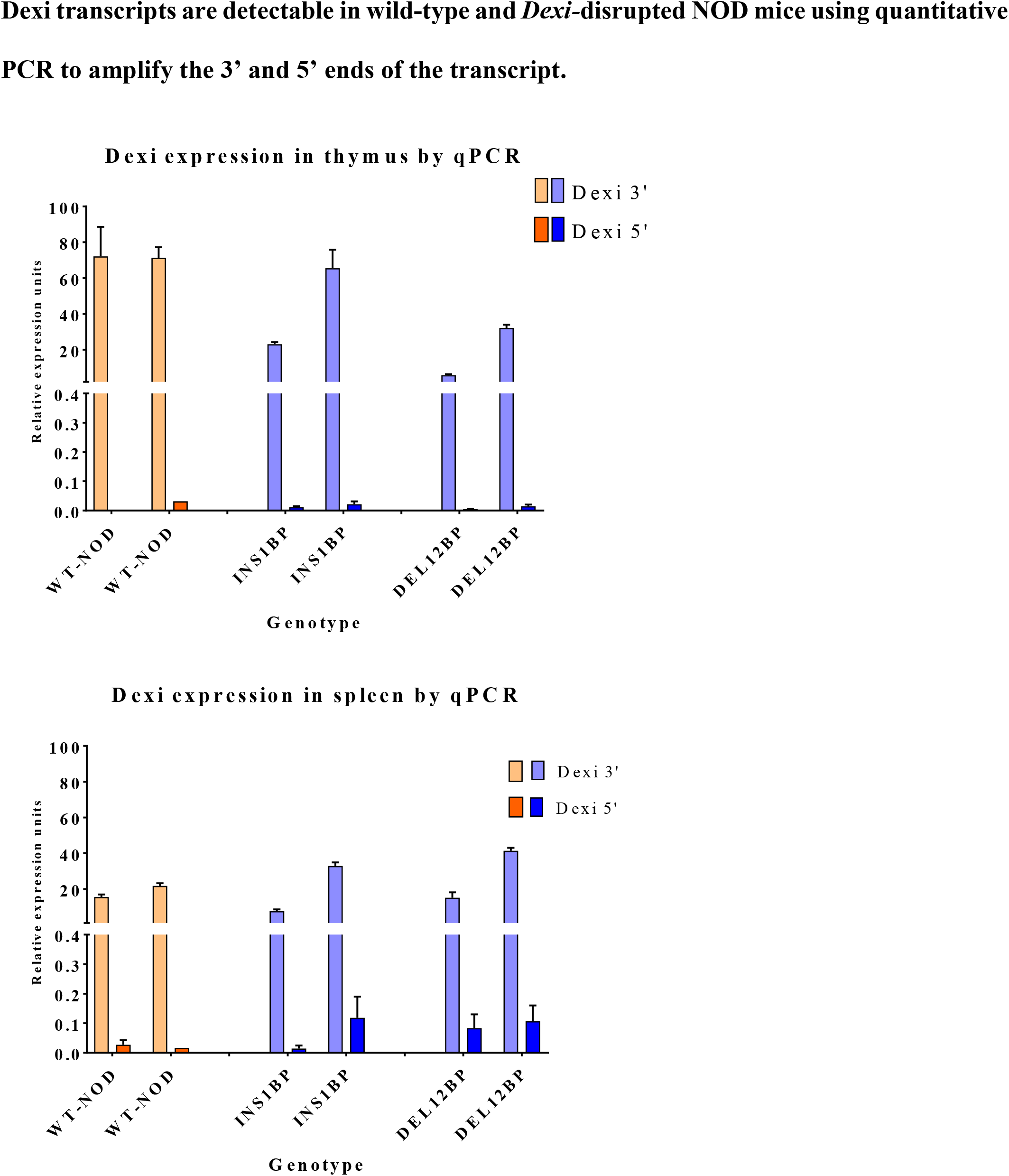
Dexi transcripts were detected in cDNA prepared from thymus and spleen using qPCR directed towards the 3’ and the 5’ end of the Dexi transcript in age-matched female wild-type (orange) NOD (n=2), homozygous *Dexi-*disrupted (blue) NOD *INS1BP* (n=2) and homozygous *Dexi-*disrupted (blue) NOD *DEL12BP* (n=2) mice. Data are shown for individual mice as mean and SD of triplicate assays using beta actin as a housekeeping gene.

**Supplementary Figure S3:**
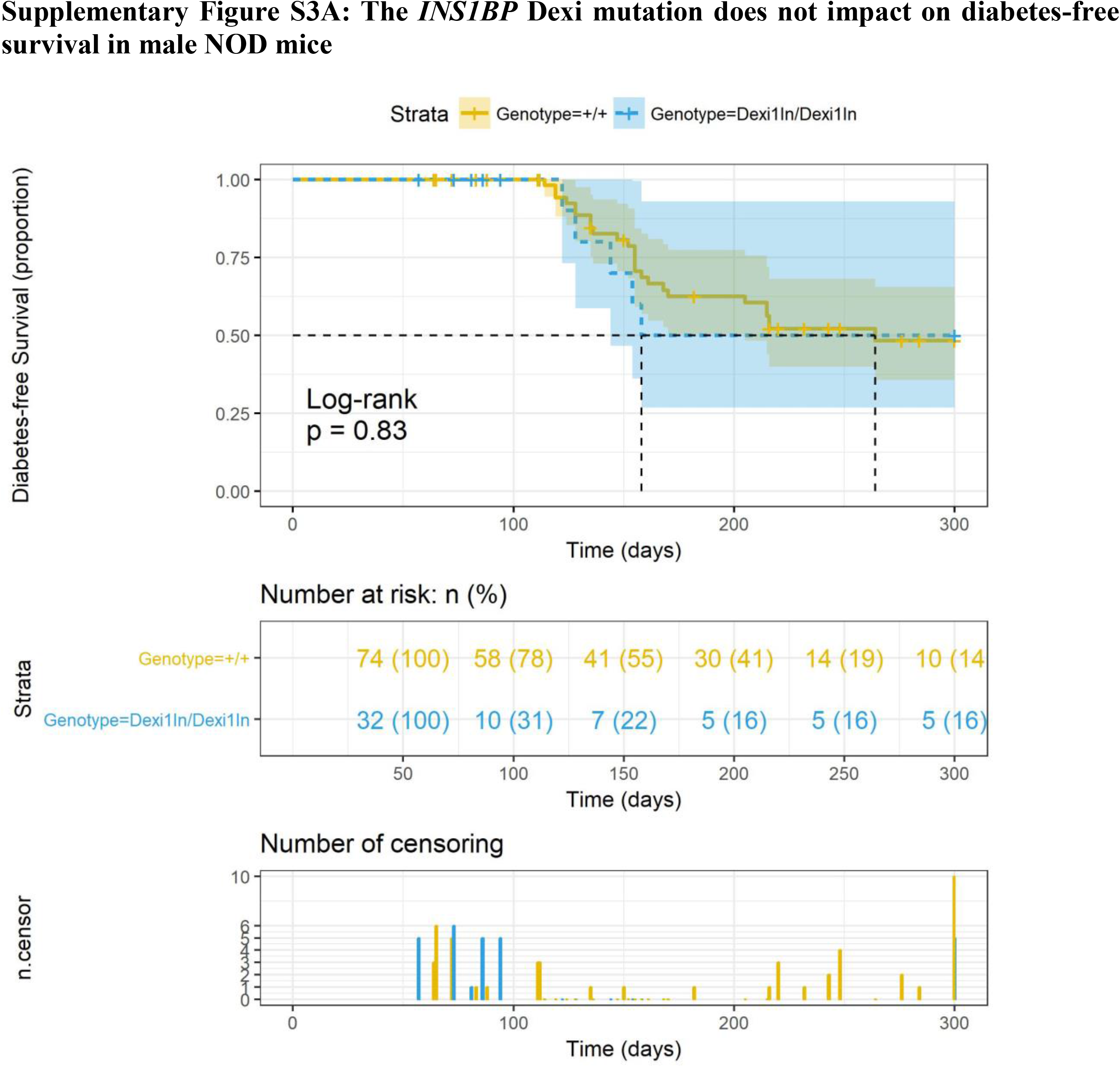
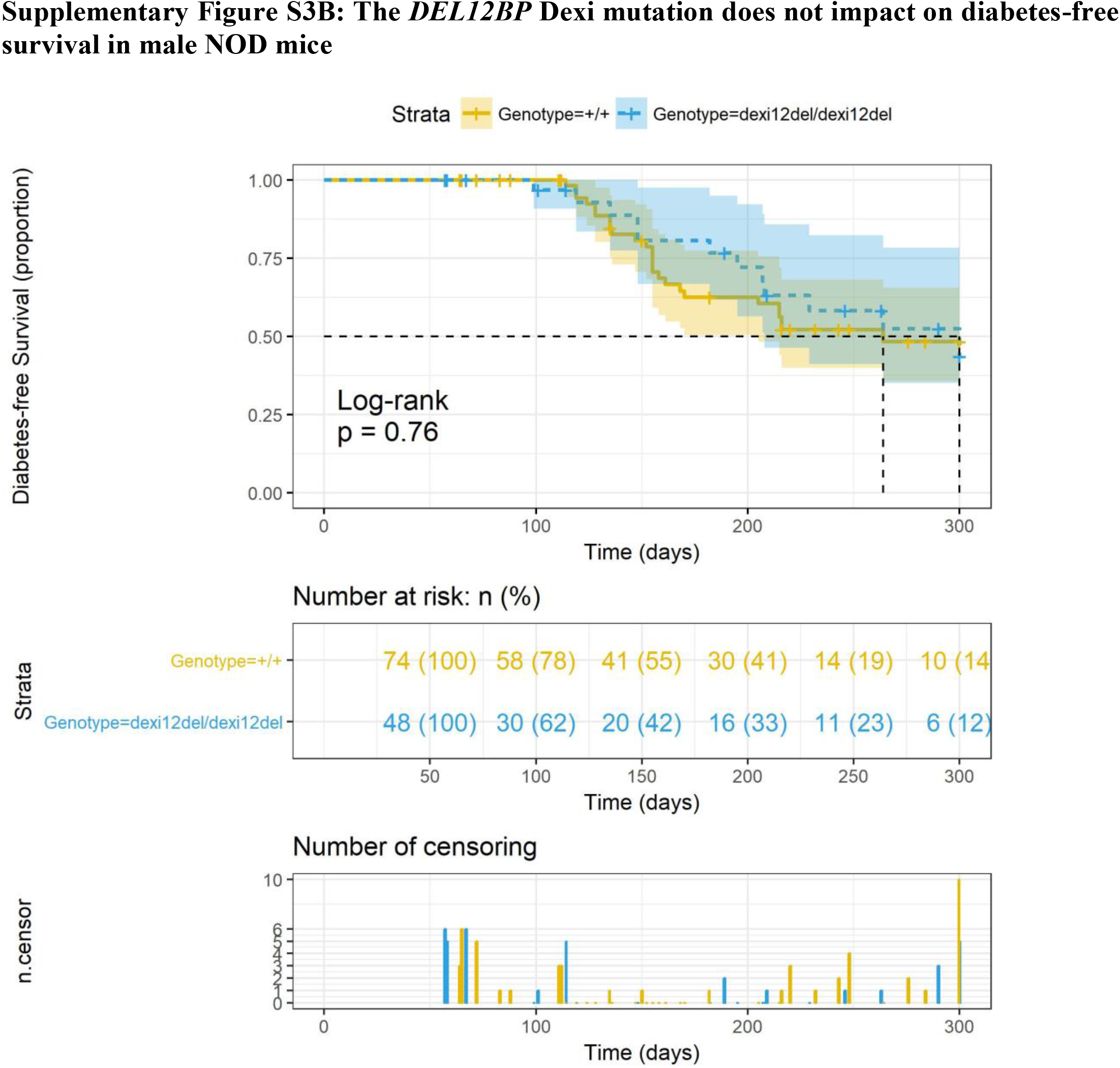
The presence of a homozygous *INS1BP* (A) or *DEL12BP* (B) *Dexi* mutation does not affect diabetes-free survival in male NOD mice. **Top panel for each line:** Kaplan-Meier diabetes-free survival curve with log-rank p-value for wild-type NOD mice (n=74) represented as yellow-orange (+/+). *Dexi*-disrupted mice survival curves are represented in blue, for homozygous *INS1BP* mice (n = 32; labelled as Dexi1in/Dexi1in) and *DEL12BP* mice (n=48); labelled as dexi12del/dexi12del). 95% Confidence Intervals are shown for each genotype as translucent shaded regions surrounding the genotype line. **Mid panel:** Numbers of animals at risk over time. **Bottom panel:** Bar plot of censorship data for each line.

**Supplementary Figure S4.**
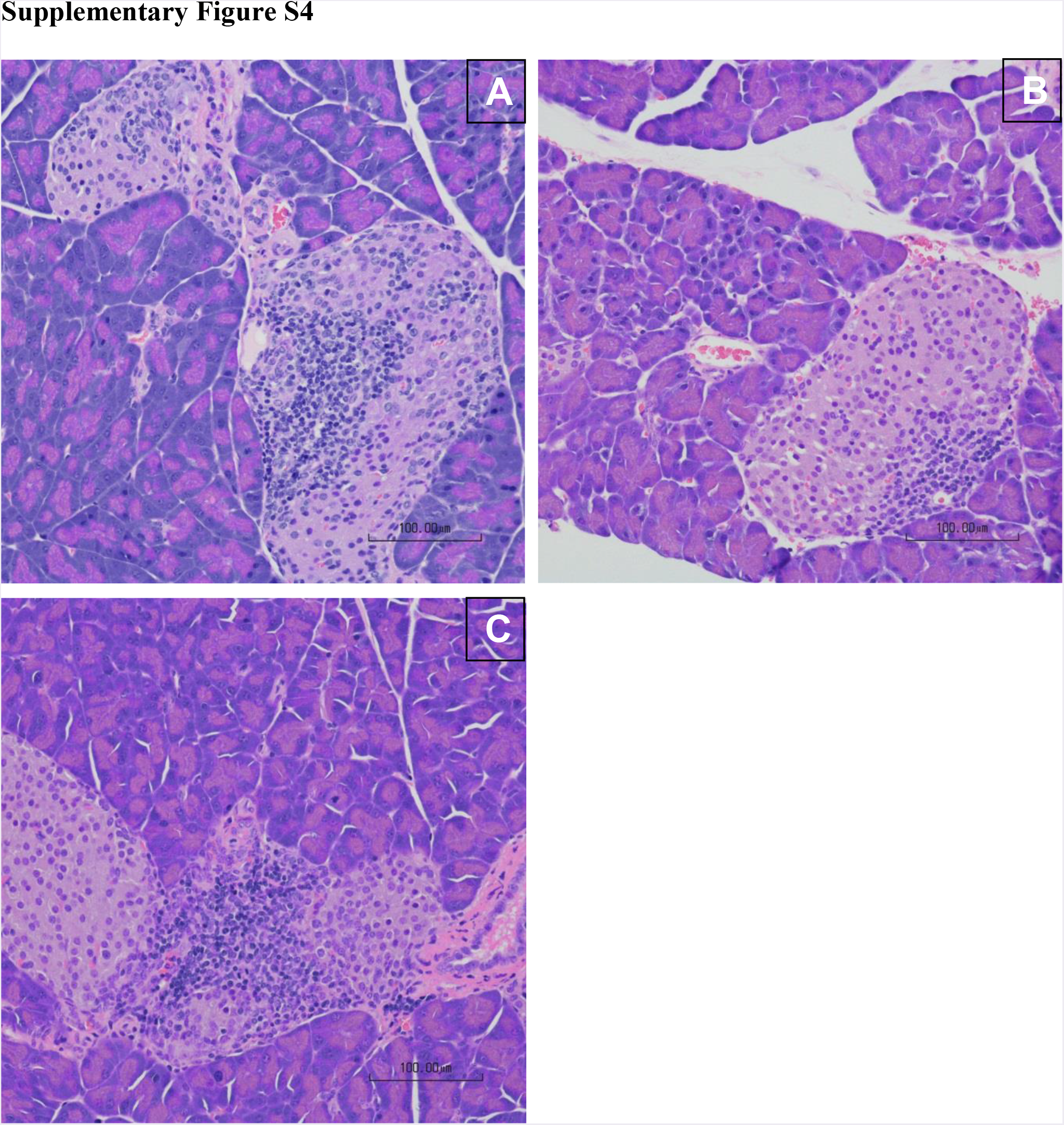
Multifocal pancreatic islets exhibit infiltration of variable numbers of lymphocytes in both NOD-WT control mice (A) and Dexi-disrupted NOD mice (B – *INS1BP-* NOD; C – DEL12BP-NOD). Haematoxylin and eosin stain. Scale bar indicates 100 microns.

**Supplementary Figure S5:**
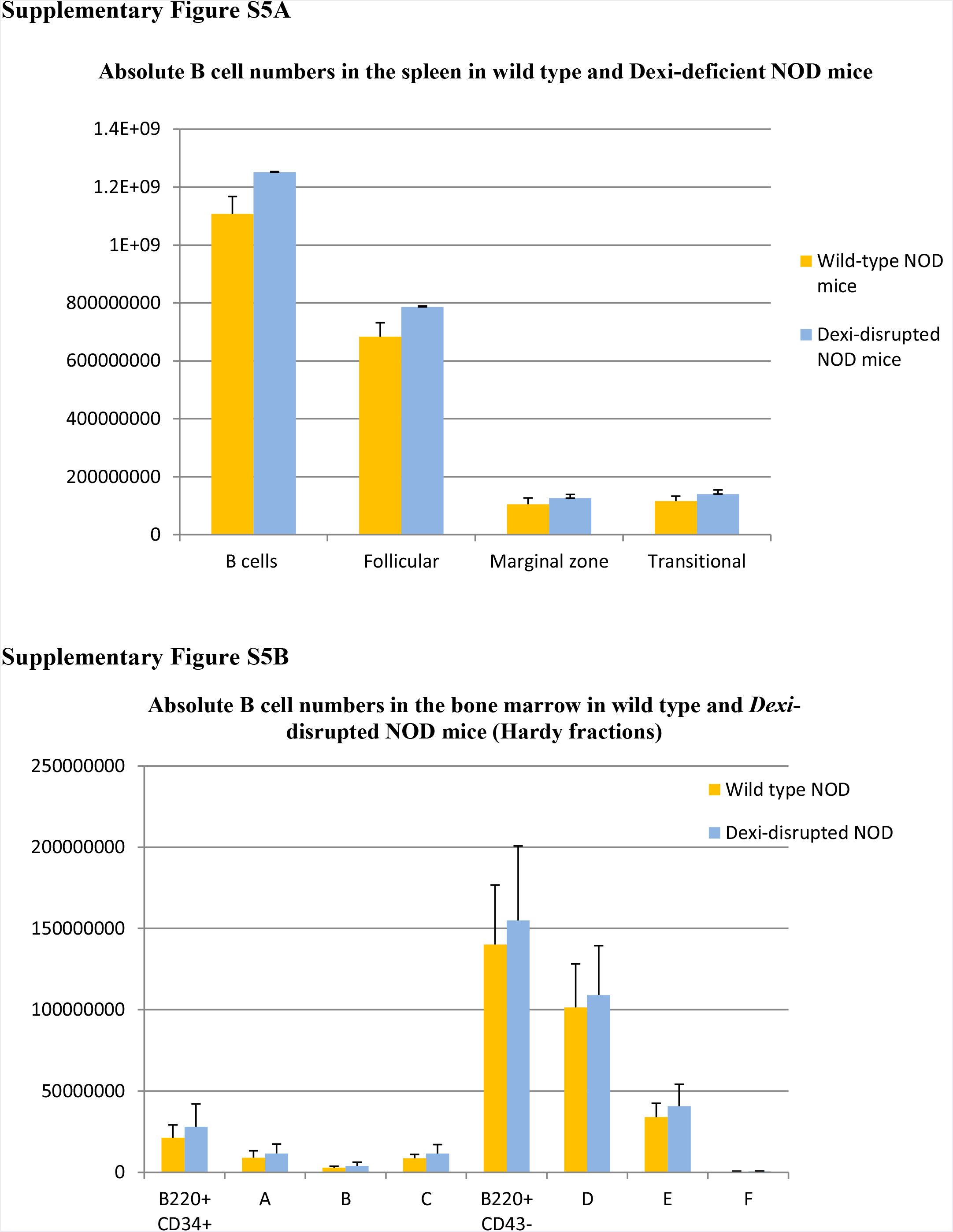
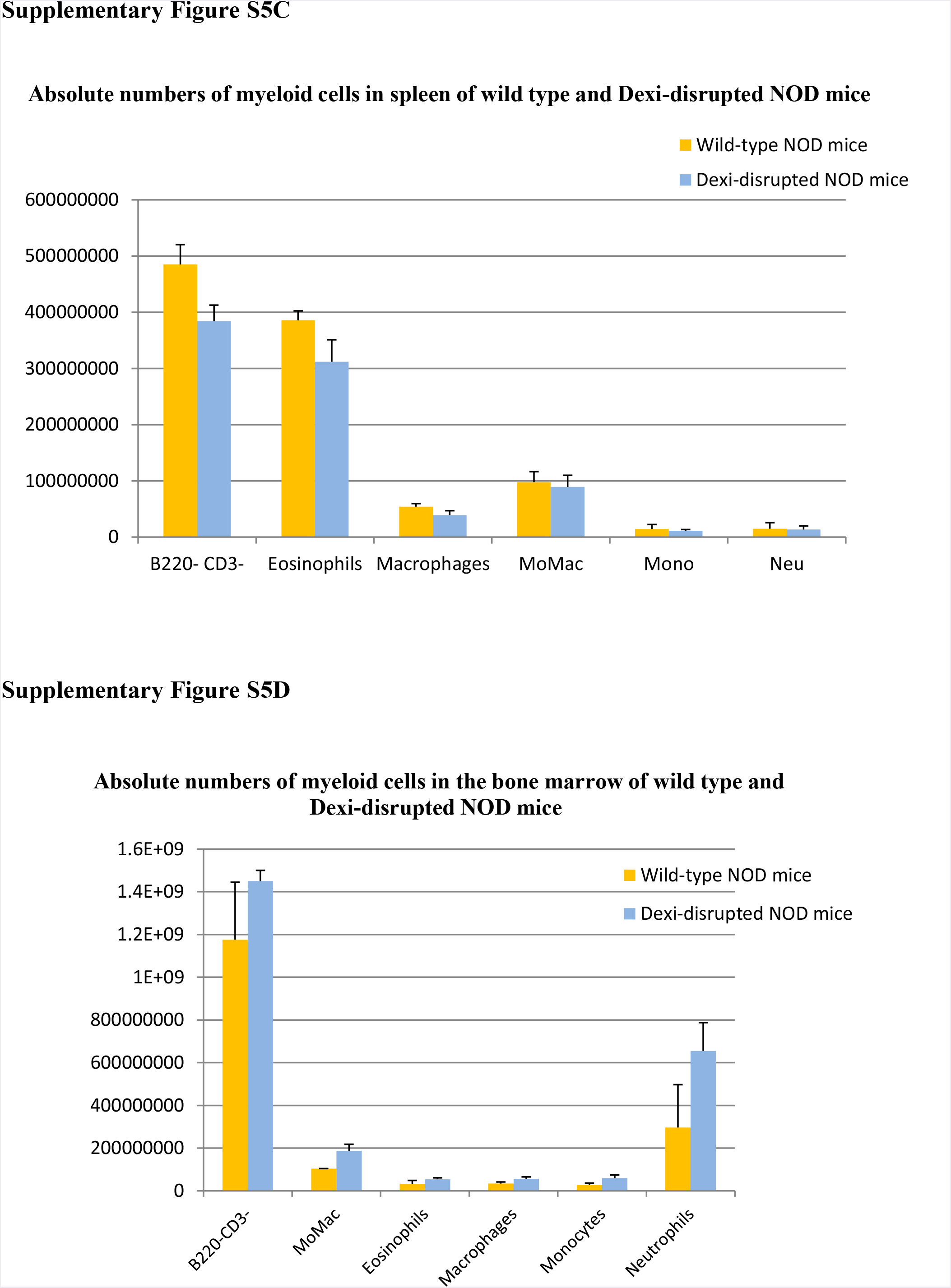
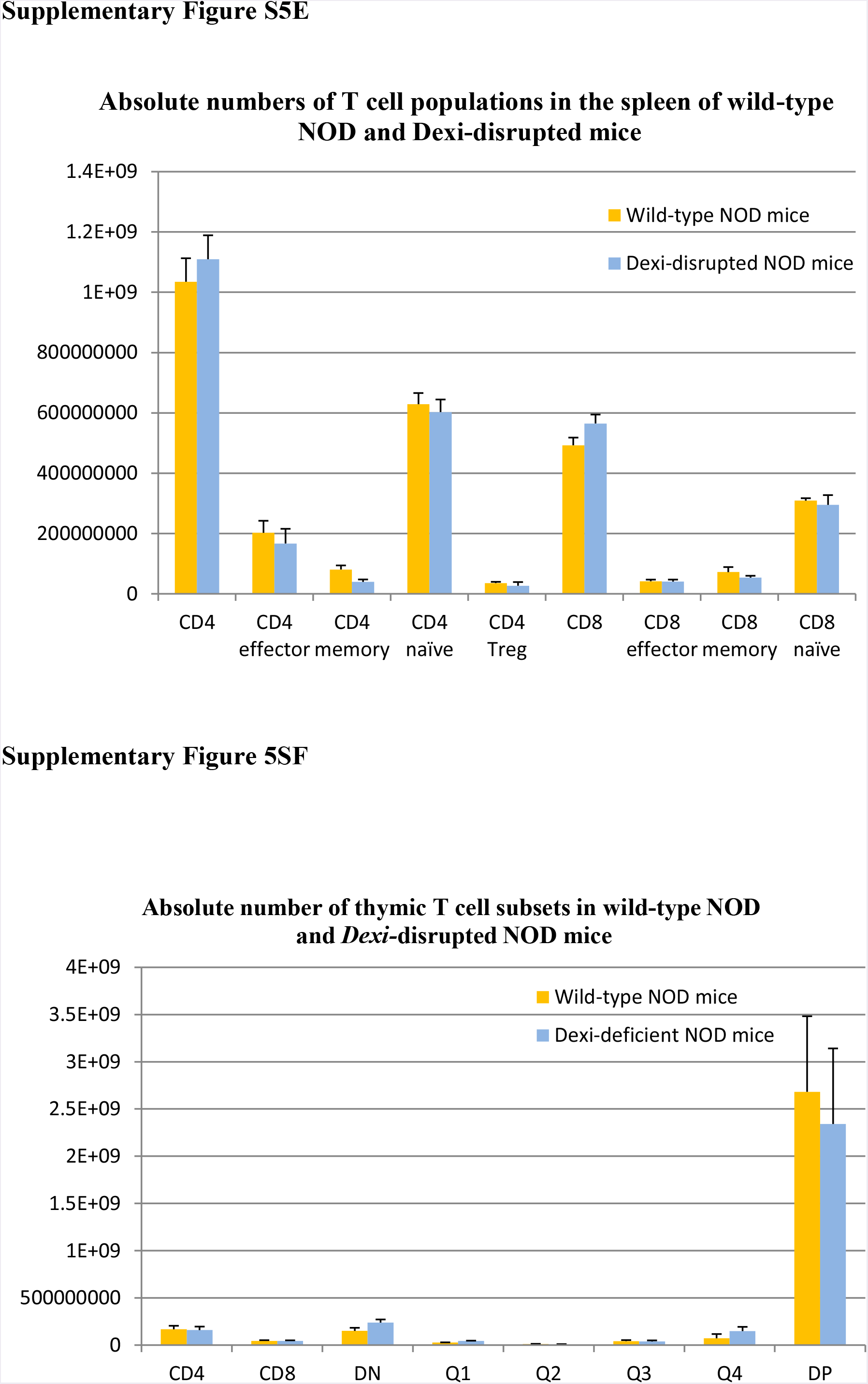
*Dexi*-disrupted mice (n=2) show no change in resting cellular immunophenotype compared to wild-type NOD mice (n=2) using a panel of monoclonal antibodies and flow cytometry. Data shown are representative of more than one experiment. See Materials and methods for gating strategies. A. Splenic B cell subsets B. Bone marrow B cell subsets C. Splenic myeloid subsets D. Bone marrow myeloid subsets E. Splenic T cells F. Thymic T cells (DN – double negative; DP double positive).

**Supplementary Figure S7:**
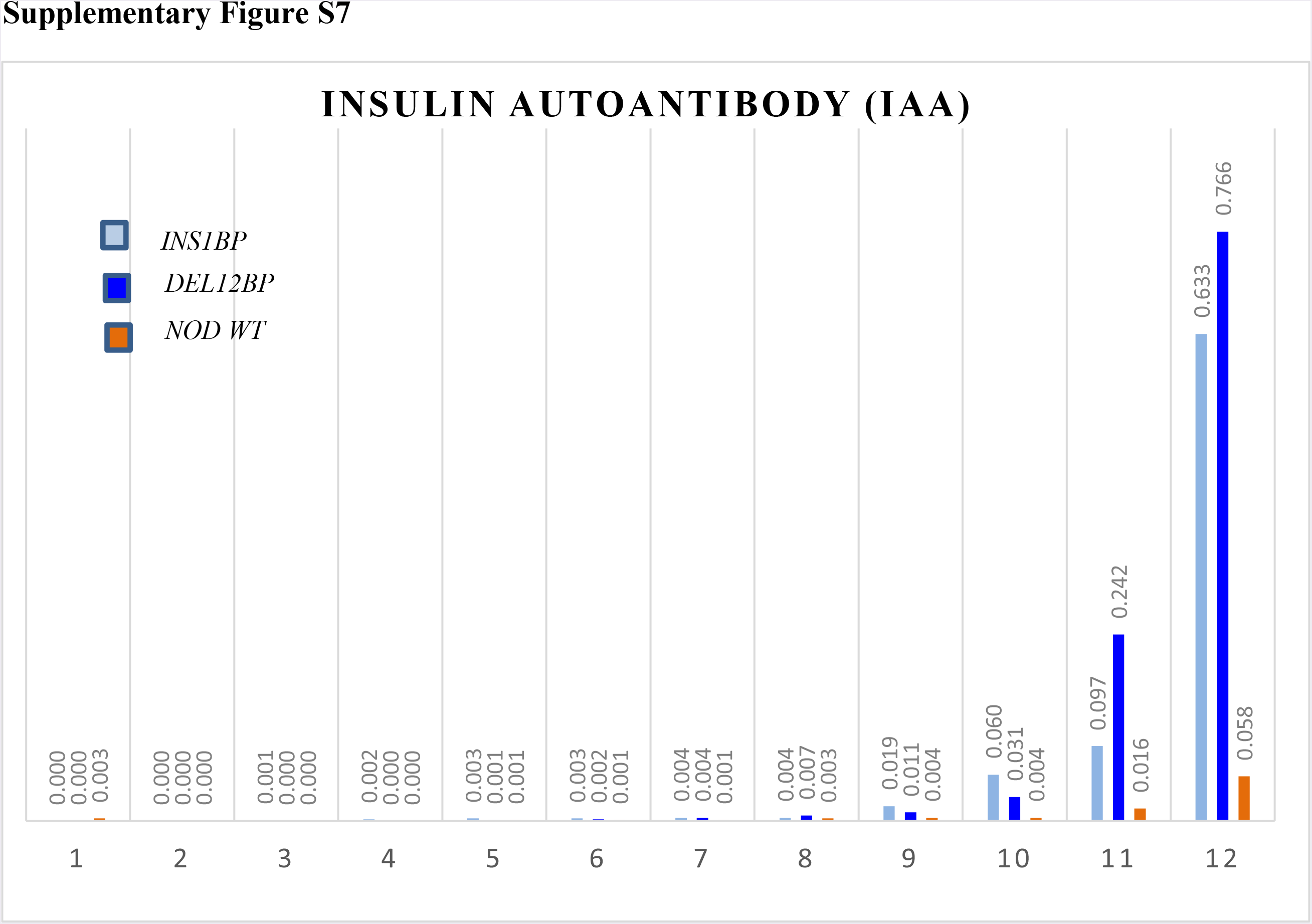
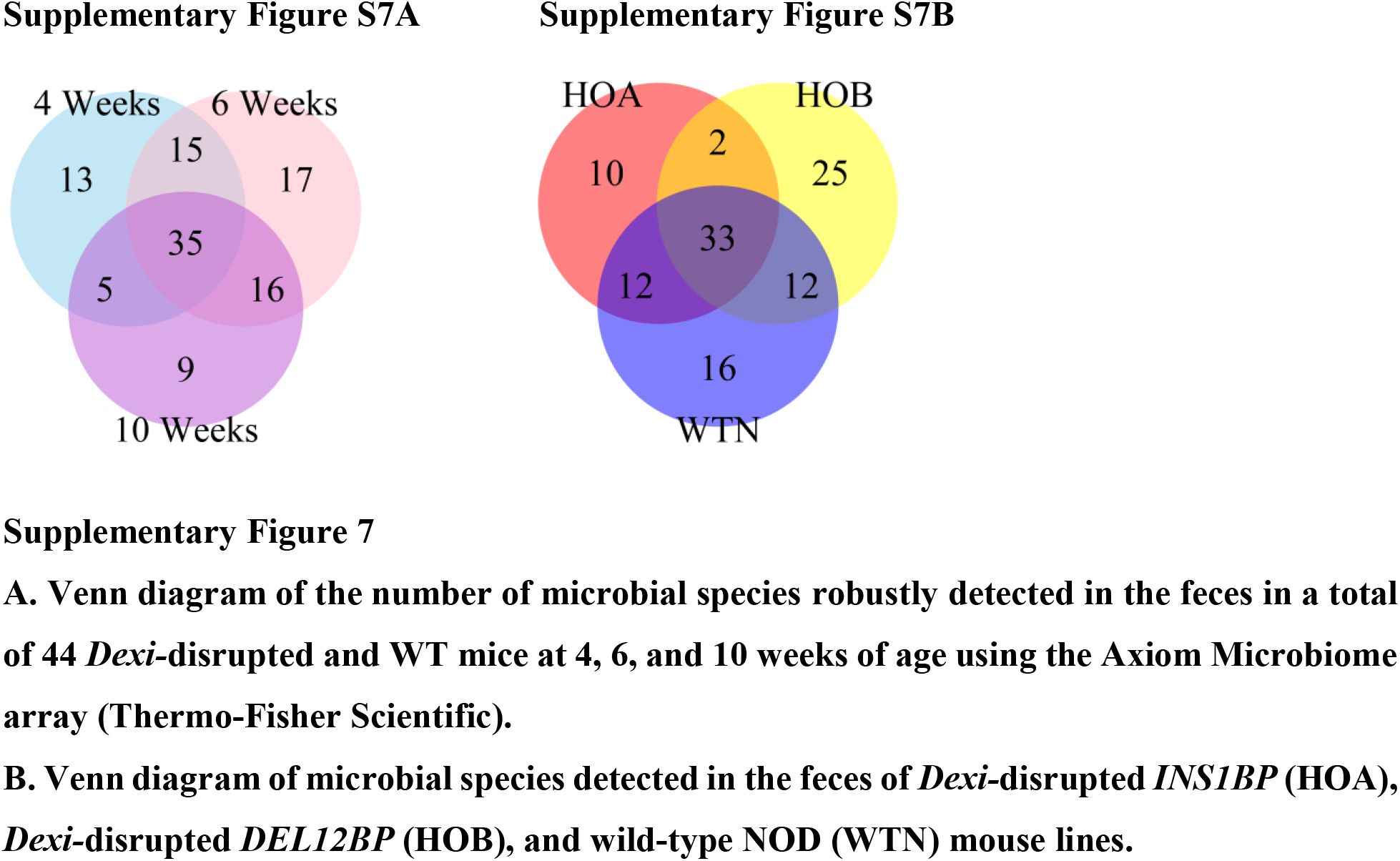
Insulin autoantibodies (IAA) were measured in the serum of 12 age-matched female mice of each genotype (NOD wild-type – NOD WT; Dexi-disrupted *INS1BP*; Dexi-disrupted *DEL12BP*) using a radioimmunoassay (at the Barbara Davis Center for Childhood Diabetes, USA). The result from each mouse (1-12) is shown individually, in ascending order of IAA. The cut-off for positivity was 0.01. Two out of 12 WT-NOD showed positivity for IAA compared to 4 out of 12 *INS1BP* mice and 4 out of 12 *DEL12BP* Dexi-disrupted mice.

**Supplementary Figure S8A.**
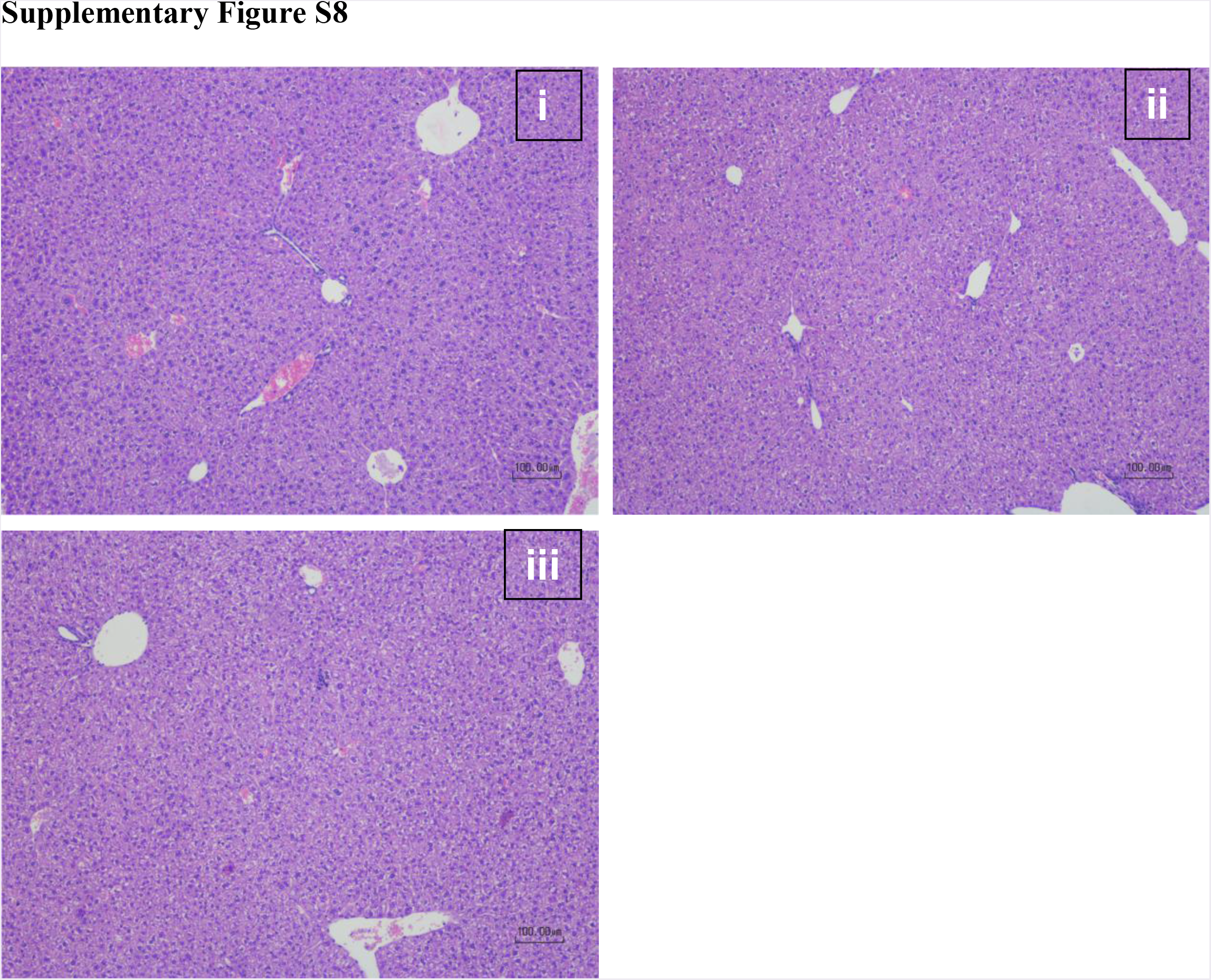
There is no histologically appreciable difference in the liver morphology of wild-type NOD mice (i) and *Dexi-*disrupted NOD mice (ii – *INS1BP* and iii –*DEL12BP*). Haematoxylin and eosin stain. Scale bar indicates 100 microns.

**Supplementary Figure 9:**
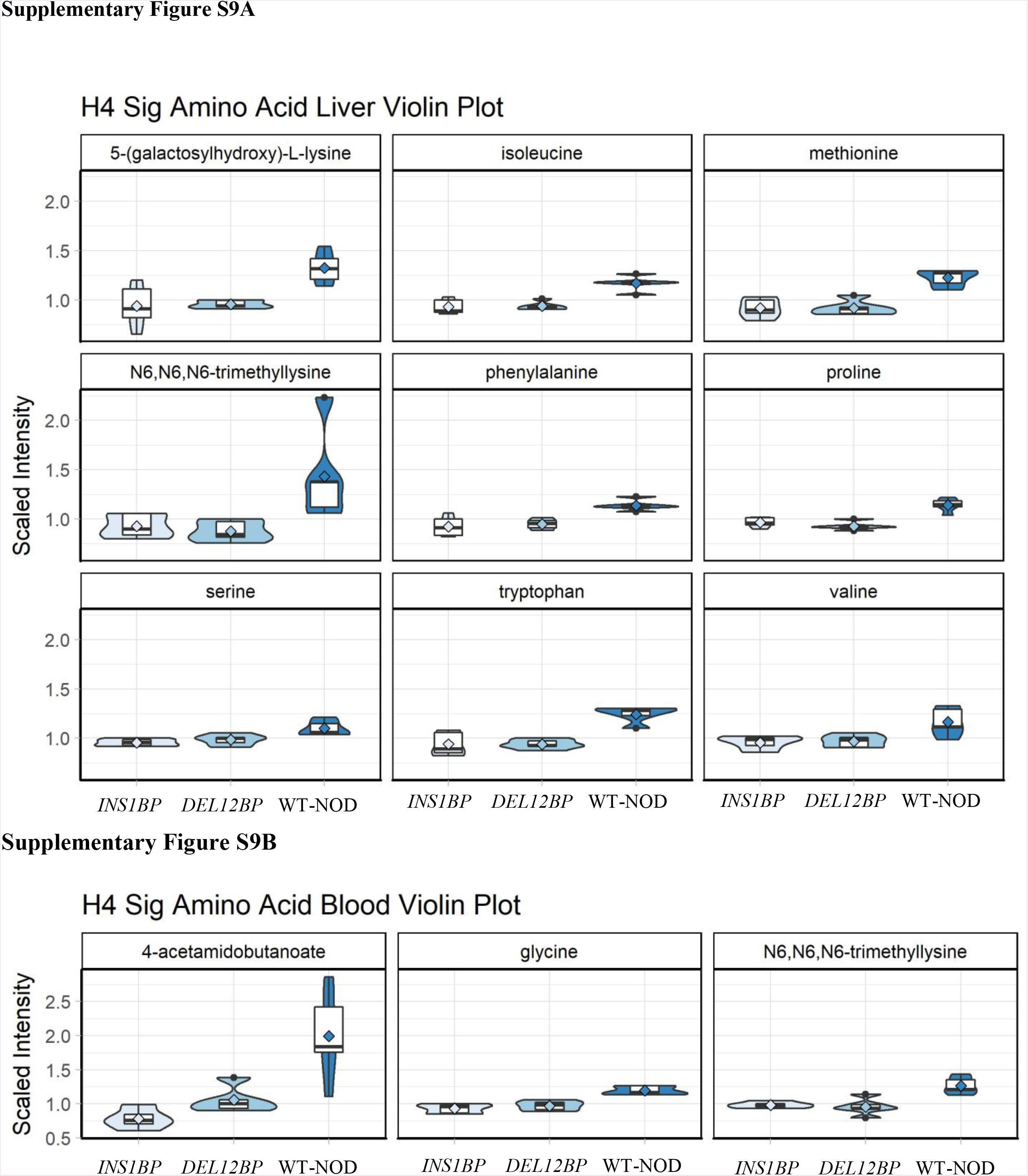
Biochemical scaled intensity violin plots of significantly different amino acids in liver (A) and blood (B). Data are shown from *INS1BP* and *DEL12BP* homozygote mice compared to wild-type (WT) NOD mice at 6 weeks of age.

**Supplementary Figure 10.**
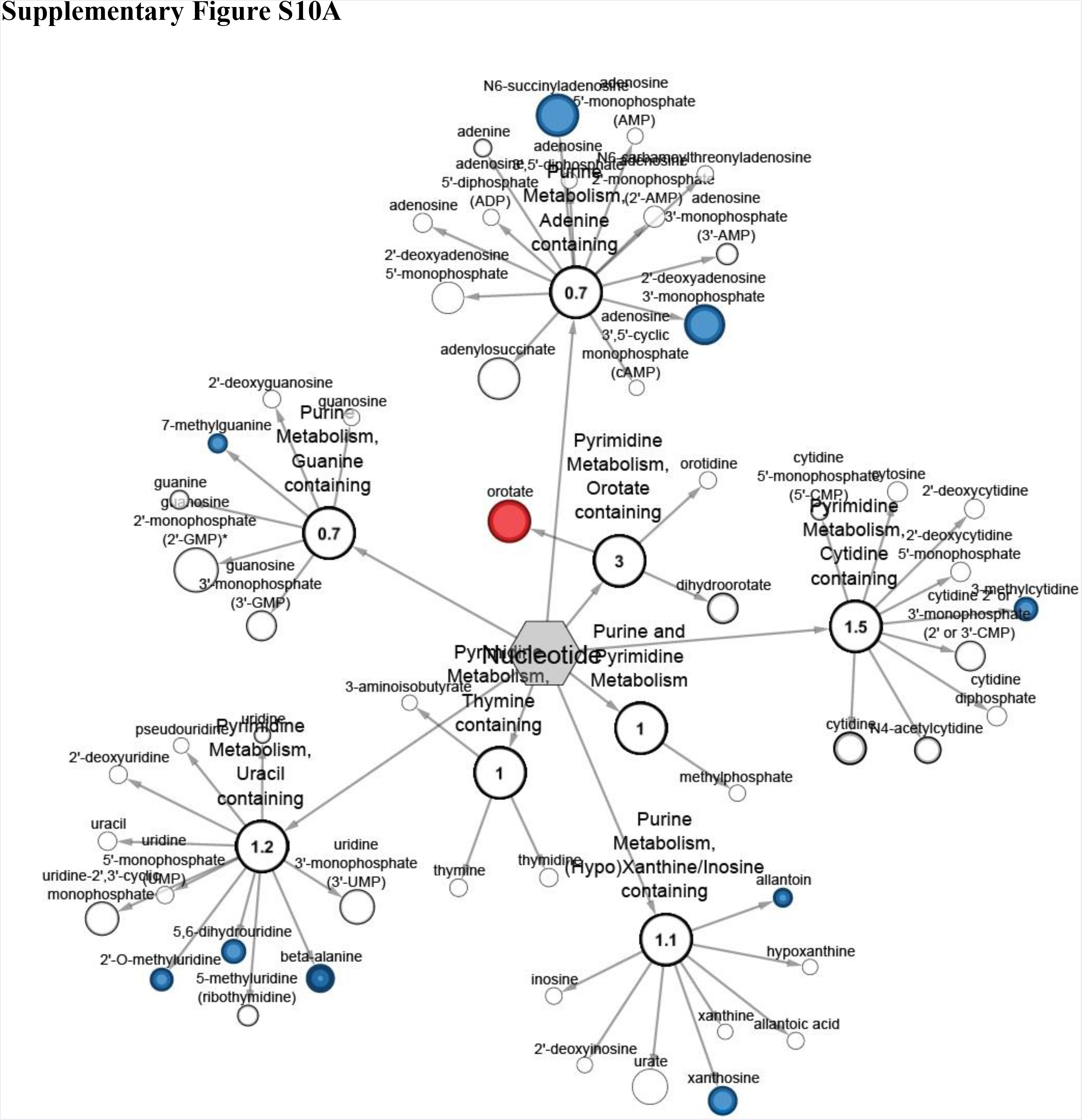
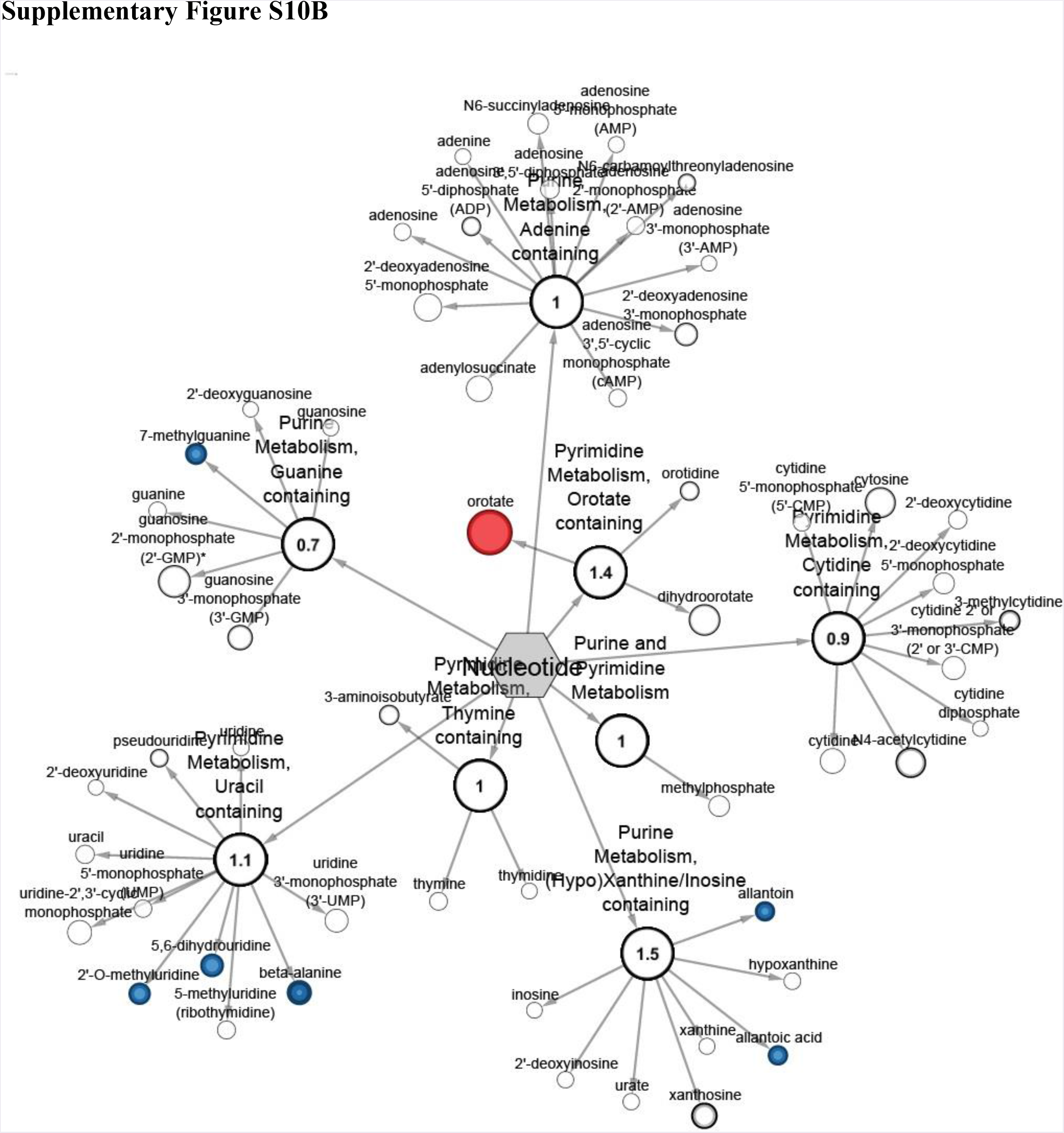
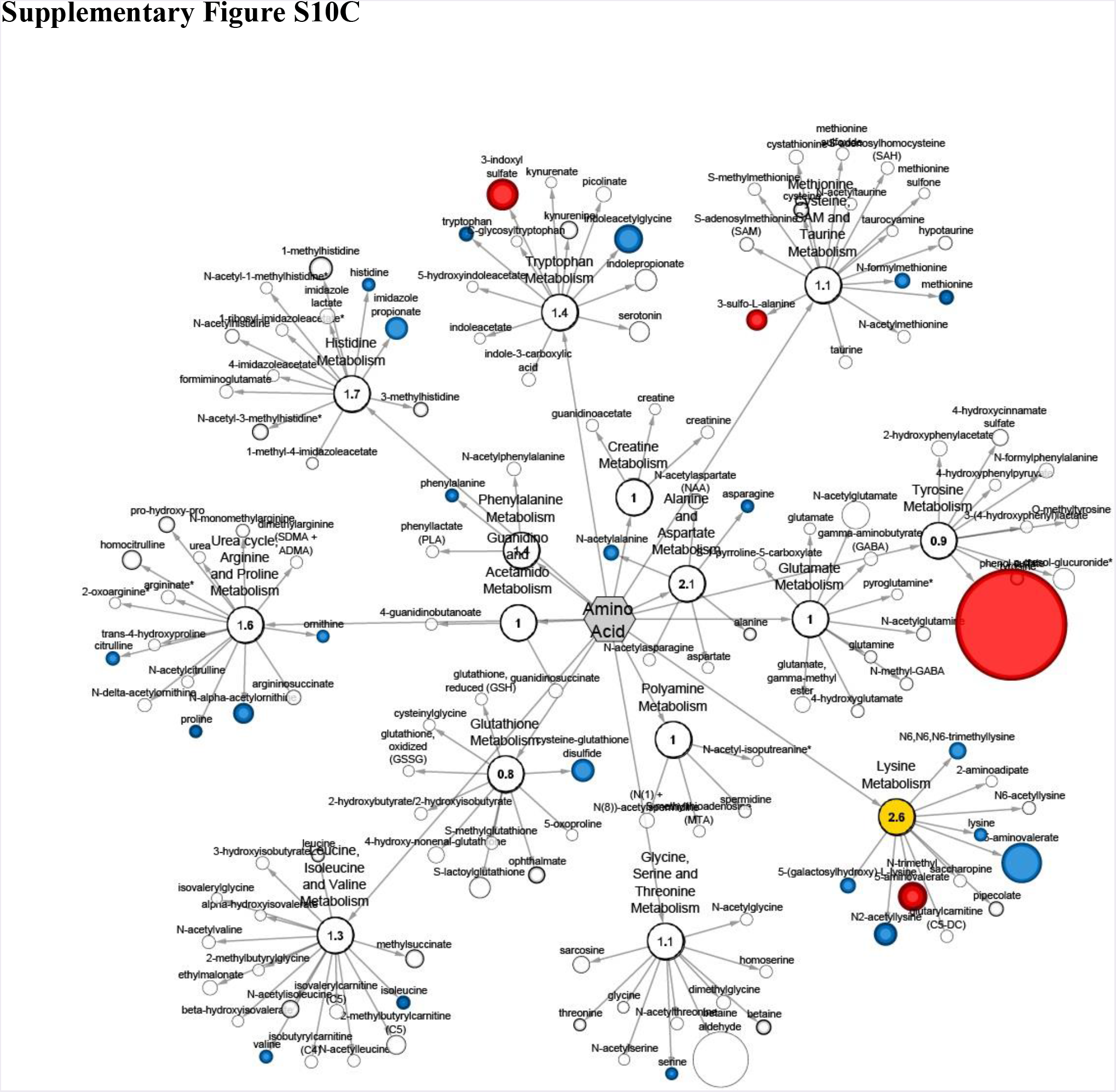
**A**. Cytoscape network for mouse liver comparing *INS1BP* **B.** mouse liver comparing *DEXL12BP* to wild-type NOD mice (Nucleotide pathway) **C**. mouse liver comparing *DEXL12BP* to wild-type NOD mice (Amino acid pathway) Shaded nodes correspond to significantly different metabolites (blue for reduced in *Dexi*-disrupted mice; red for increased in *Dexi*-disrupted mice; yellow for a significant difference in the whole pathway). Dark red or blue represent change at p < 0.05 and lighter colored nodes correspond to trending threshold p < 0.1. White nodes correspond to detected compounds with no significant change.

**Supplementary Table S1.**
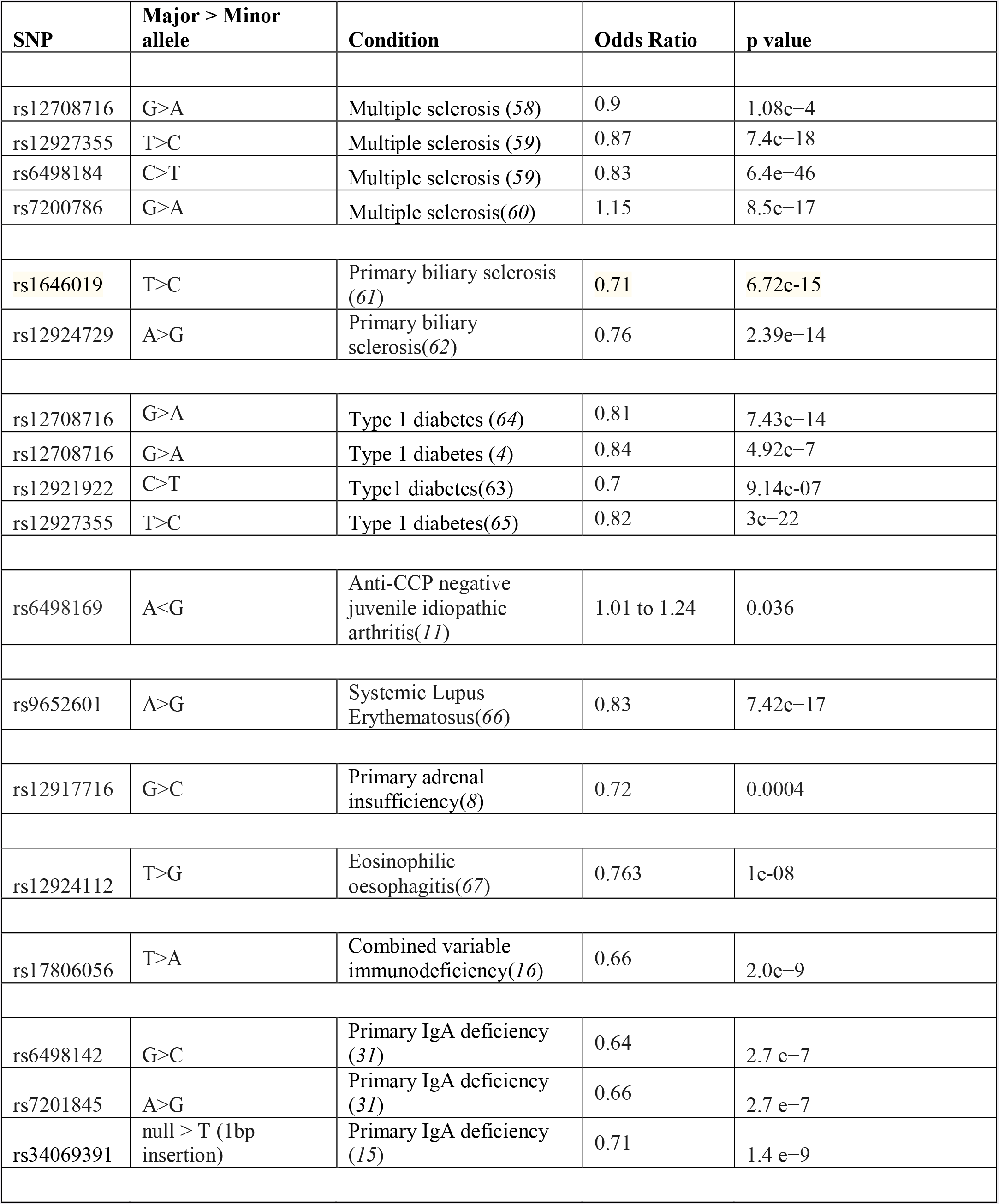
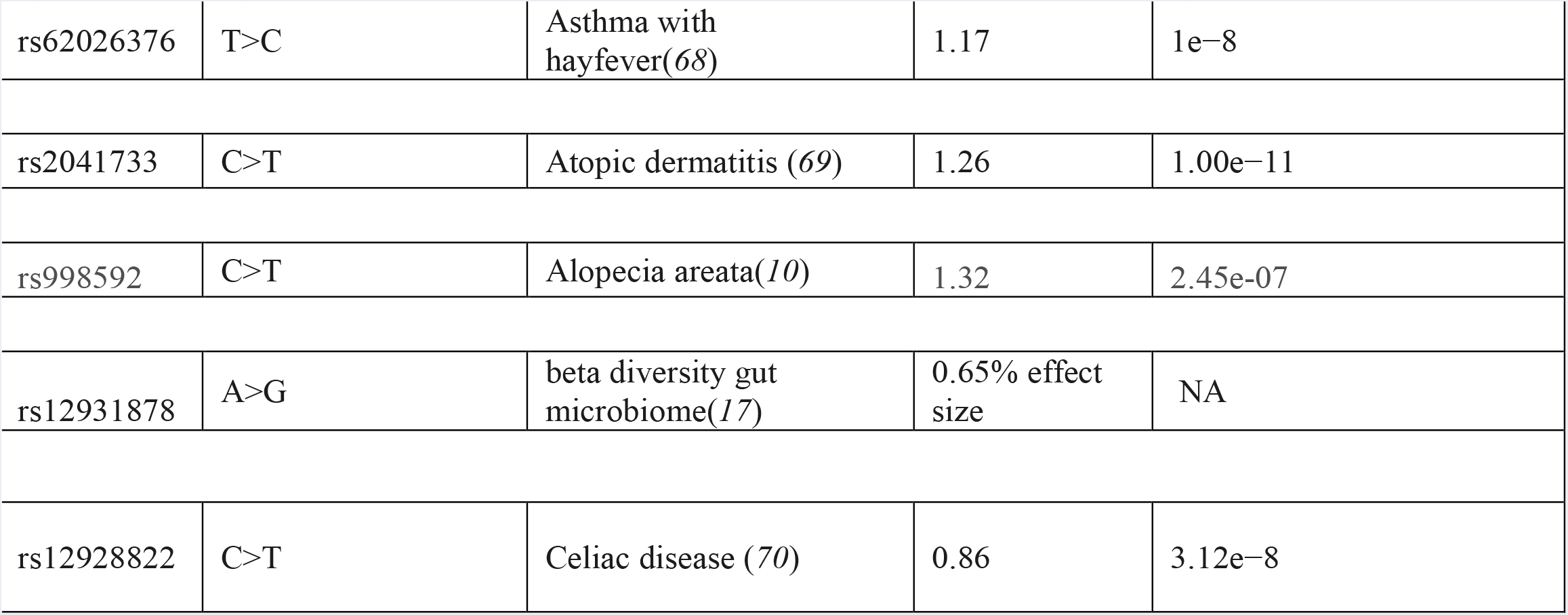
Examples of published SNPs in 16p13.13 region associated with immune-related conditions.

| SNP | Major > Minor allele | Condition | Odds Ratio | p value |
| --- | --- | --- | --- | --- |
| rs12708716 | G>A | Multiple sclerosis (58) | 0.9 | 1.08e-4 |
| rs12927355 | T>C | Multiple sclerosis (59) | 0.87 | 7.4e-18 |
| rs6498184 | C>T | Multiple sclerosis (59) | 0.83 | 6.4e-46 |
| rs7200786 | G>A | Multiple sclerosis(60) | 1.15 | 8.5e-17 |
| rs1646019 | T>C | Primary biliary sclerosis (61) | 0.71 | 6.72e-15 |
| rs12924729 | A>G | Primary biliary sclerosis(62) | 0.76 | 2.39e-14 |
| rs12708716 | G>A | Type 1 diabetes (64) | 0.81 | 7.43e-14 |
| rs12708716 | G>A | Type 1 diabetes (4) | 0.84 | 4.92e-7 |
| rs12921922 | C>T | Type1 diabetes(63) | 0.7 | 9.14e-07 |
| rs12927355 | T>C | Type 1 diabetes(65) | 0.82 | 3e-22 |
| rs6498169 | A<G | Anti-CCP negative juvenile idiopathic arthritis(11) | 1.01 to 1.24 | 0.036 |
| rs9652601 | A>G | Systemic Lupus Erythematosus(66) | 0.83 | 7.42e-17 |
| rs12917716 | G>C | Primary adrenal insufficiency(8) | 0.72 | 0.0004 |
| rs12924112 | T>G | Eosinophilic oesophagitis(67) | 0.763 | 1e-08 |
| rs17806056 | T>A | Combined variable immunodeficiency(16) | 0.66 | 2.0e-9 |
| rs6498142 | G>C | Primary IgA deficiency (31) | 0.64 | 2.7 e-7 |
| rs7201845 | A>G | Primary IgA deficiency (31) | 0.66 | 2.7 e-7 |
| rs34069391 | null > T (1bp insertion) | Primary IgA deficiency (15) | 0.71 | 1.4 e-9 |
| rs62026376 | T>C | Asthma with hayfever(68) | 1.17 | 1e-8 |
| rs2041733 | C>T | Atopic dermatitis (69) | 1.26 | 1.00e-11 |
| rs998592 | C>T | Alopecia areata(10) | 1.32 | 2.45e-07 |
| rs12931878 | A>G | beta diversity gut microbiome(17) | 0.65% effect size | NA |
| rs12928822 | C>T | Celiac disease (70) | 0.86 | 3.12e-8 |

**Supplementary Table S2.**
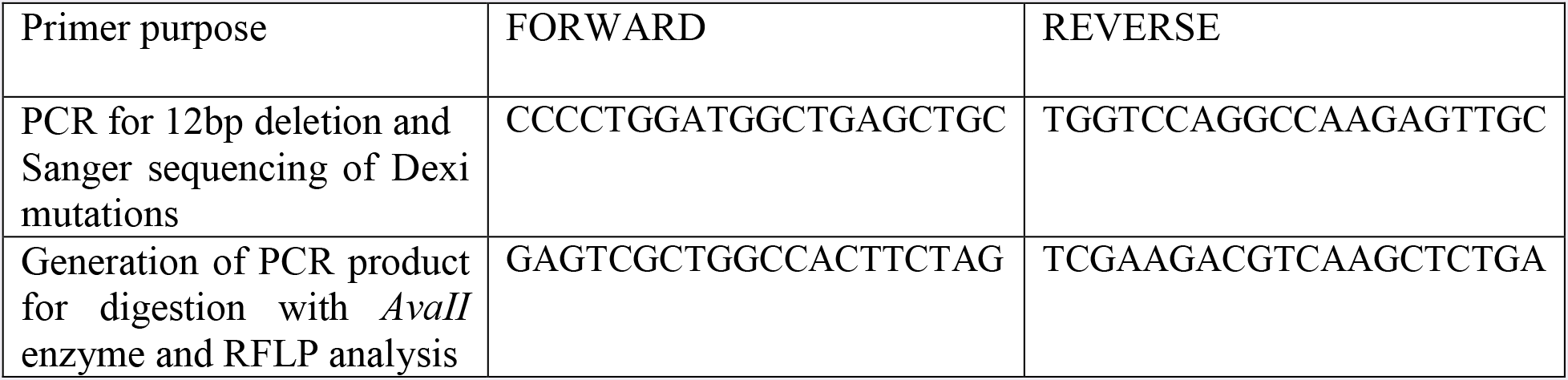
– Primers for genotyping.

**Supplementary Table S3:** Mice included in the diabetes incidence study.

| *Cohort* | *Sex* | *Mantel Haenszel p-value* | *Dexi-disrupted mice #* | *Mean Dexi-disrupted diabetes onset (days)* | *Median Dexi-disrupted diabetes onset (days)* | *WT NOD Sample #* | *Mean WT NOD diabetes onset (days)* | *Median WT NOD diabetes onset (days)* |
| --- | --- | --- | --- | --- | --- | --- | --- | --- |
| *INS1BP* | Female | **0.0254** | 37 | 110 | 113 | 79 | 153 | 129 |
| *INS1BP* | Male | 0.8284 | 32 | 122 | 86 | 74 | 172 | 155 |
| *DEL12BP* | Female | **0.0307** | 77 | 122 | 114 | 79 | 153 | 129 |
| *DEL12BP* | Male | 0.7649 | 48 | 154 | 117 | 74 | 172 | 155 |

**Supplementary Table S4:**
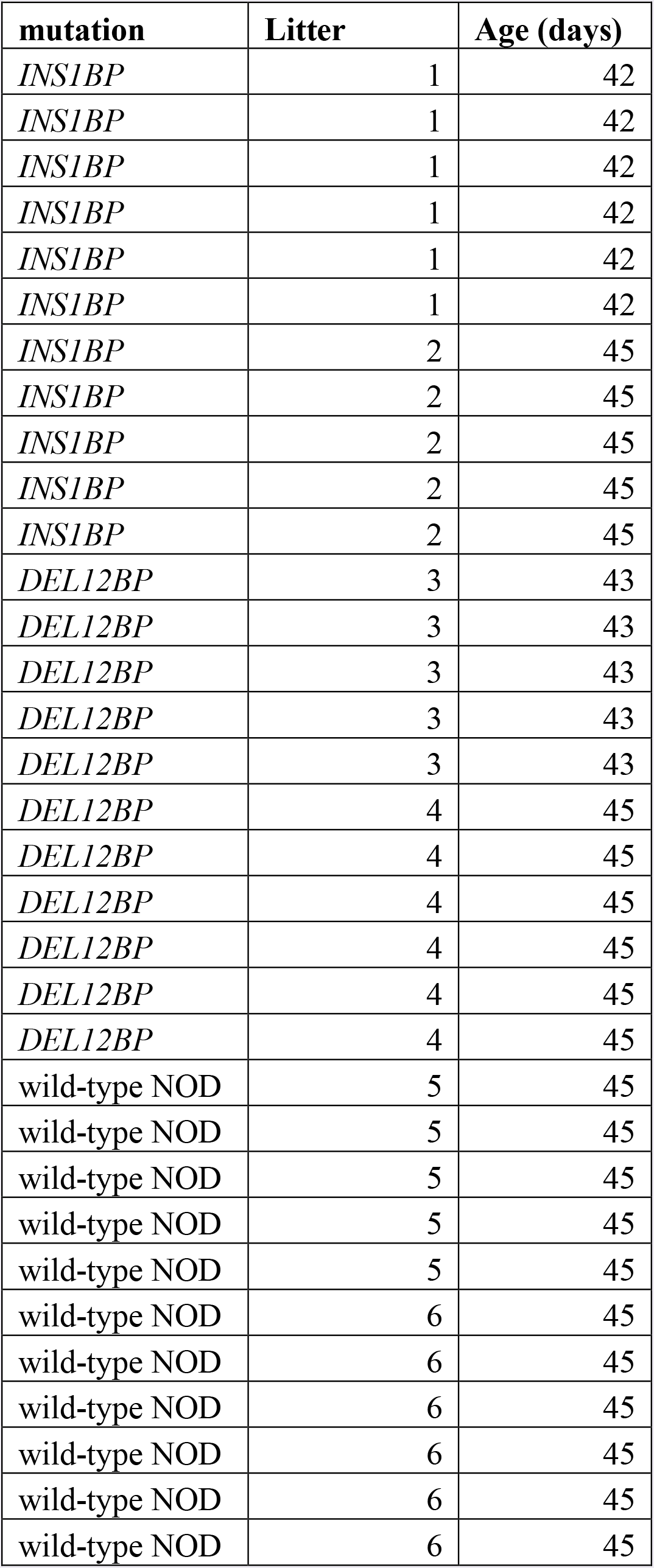
Mice included in the histopathological islet study.

**Supplementary Table S5.** Antibodies for flow cytometry.

| **B cell panel (BM)** | **B cell panel (Spleen)** | **Myeloid panel** | **T cell panel** |
| --- | --- | --- | --- |
| CD24-FITC | IgD-FITC | CD11c-FITC | CD44-FITC |
| BPI-PE | IgM-PE | CD8a-PE | CD25-PE |
| CD19-PerCP | CD19-PerCP | Ly6c-PerCP | CD8a-PerCP |
| IgM-PECy7 | CD23-PECy7 | F4/80-PECy7 |  |
| CD43-APC | MHCII-APC (I-Ad) | CD11b-APC | CD62-APC |
| B220 -AF700 | B220-AF700 | Ly6g Gr1-AF700 | CD4-AF700 |
| IgD-APC-Cy7 | | CD3e-PB | TCR$\alpha\beta$ APC-Cy7 |
|  | CD21-B605 | B220-BV605 | CD69-BV605 |

**Supplementary Table S6.**
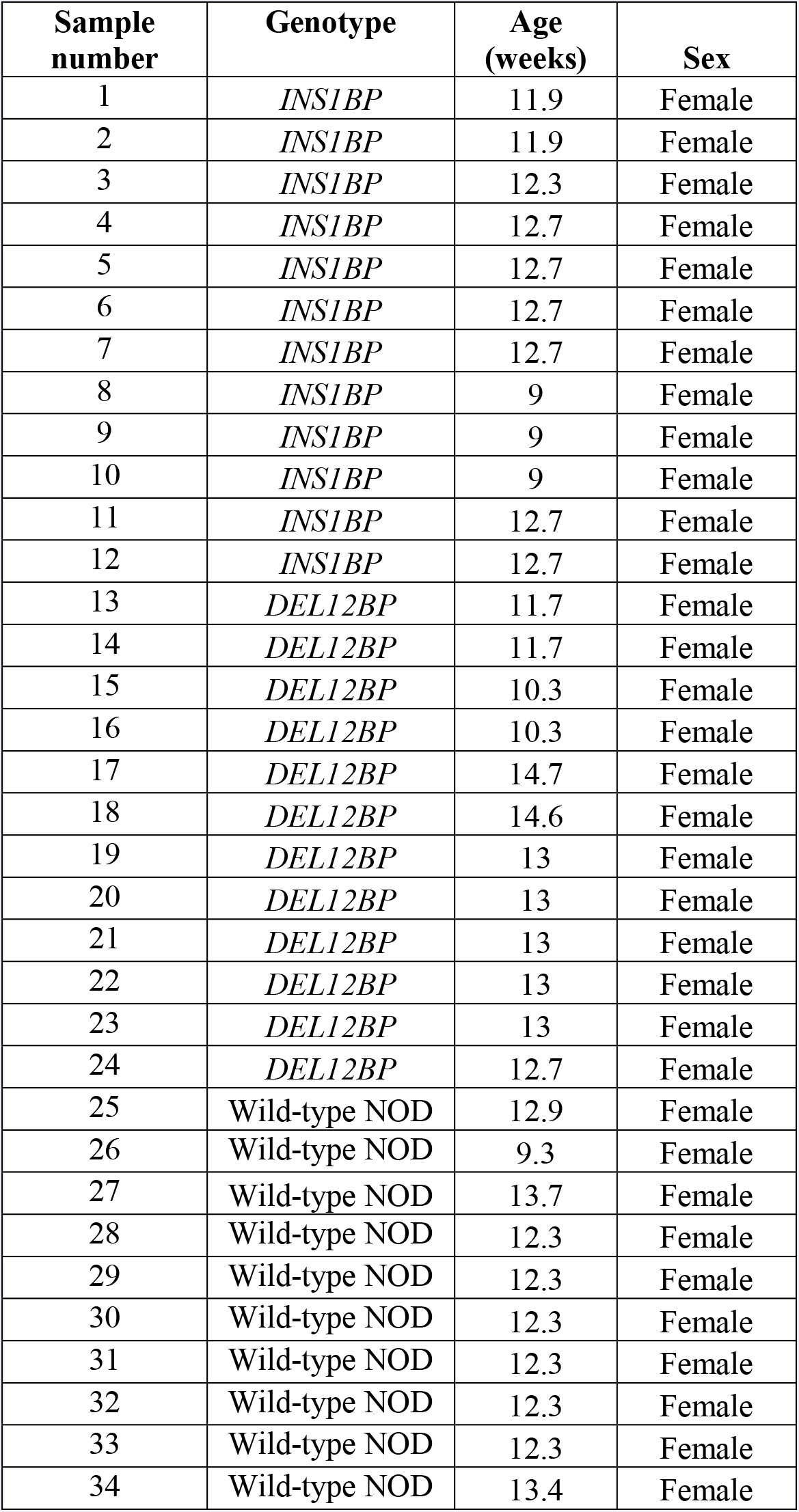
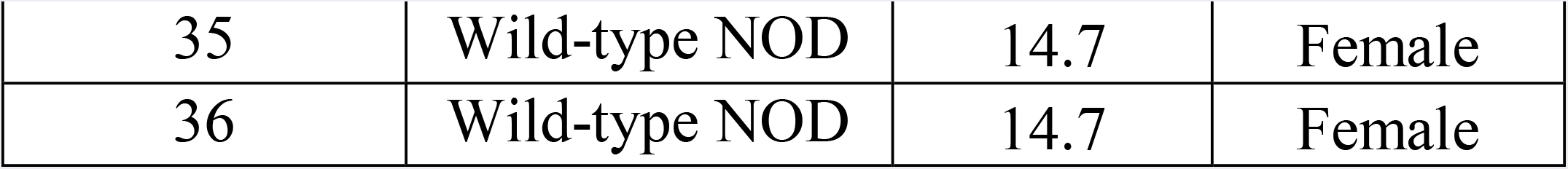
– Descriptive statistics of mice included in ELISA experiments All mice were normoglycemic and were aged between 9 and 15 weeks. *Dexi*-disrupted mice were homozygous for either the *DEL12BP* mutation or the *INS1BP* mutation on the NOD genetic background.

**Supplementary Table S7.**
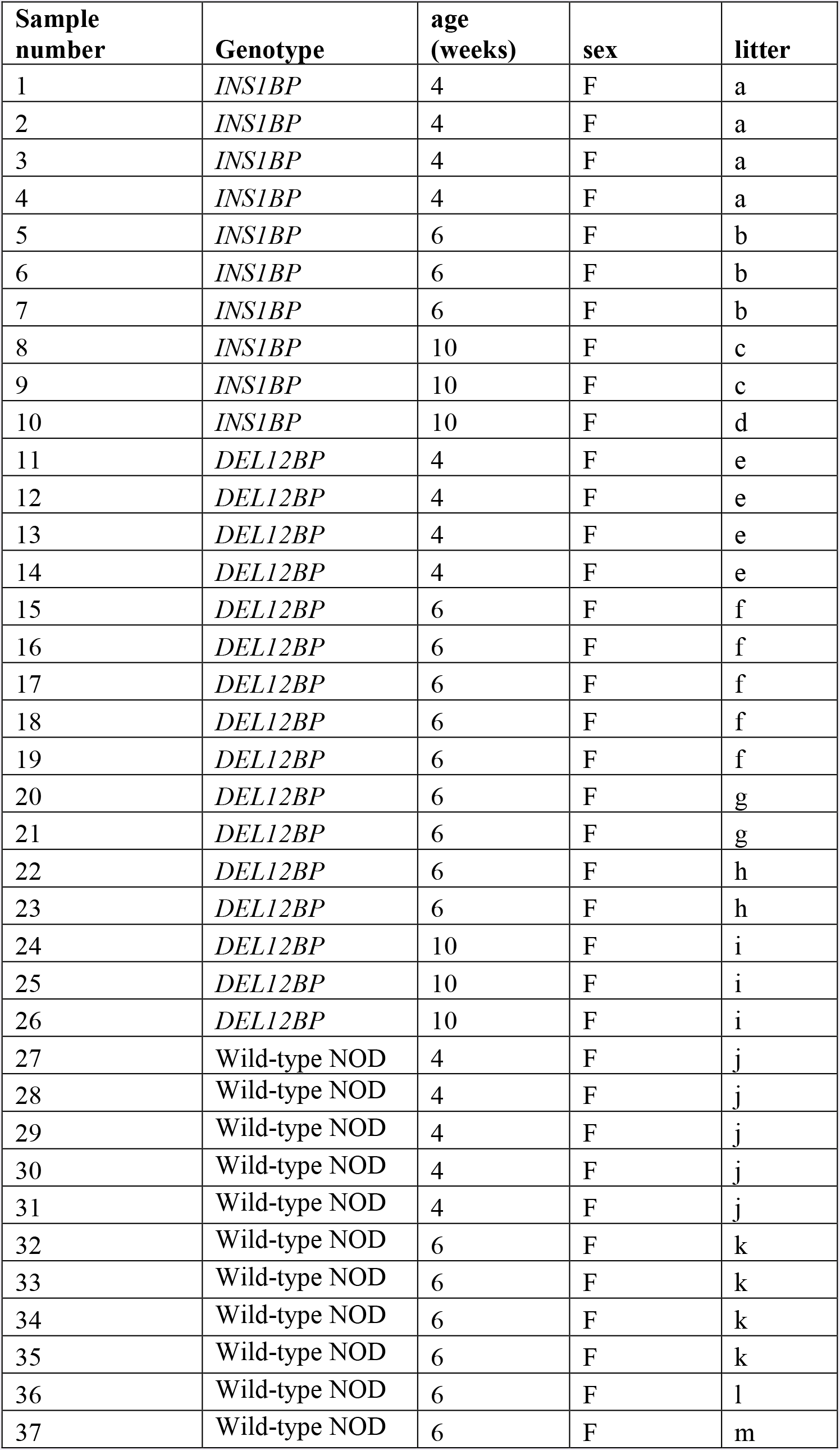
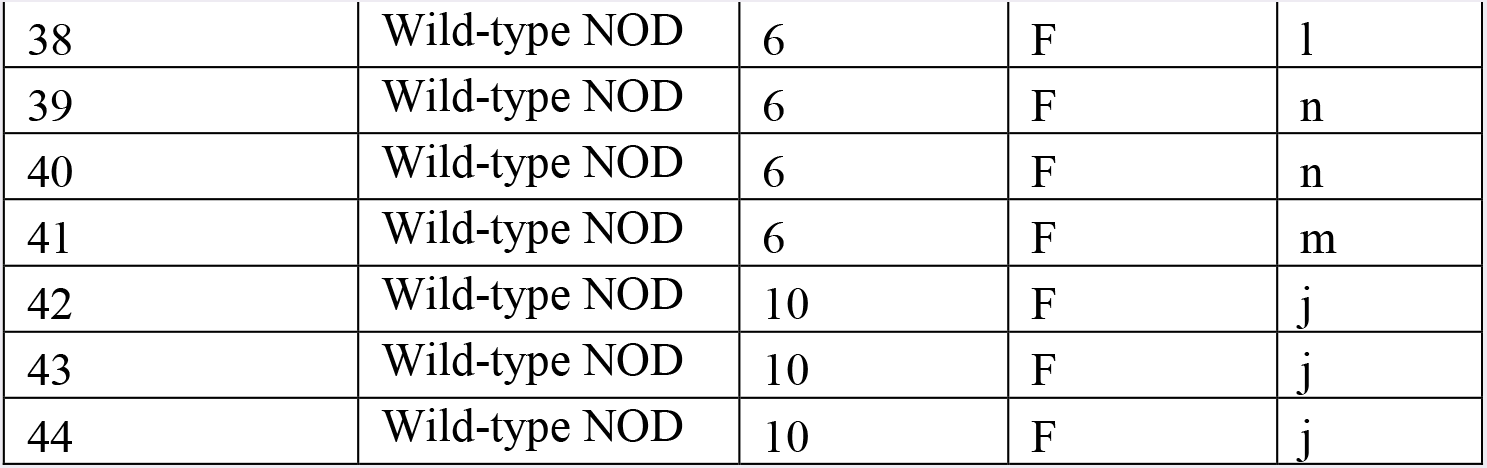
– Mice included in the microbiome study.

| **Sample number** | **Genotype** | **age (weeks)** | **sex** | **litter** |
| --- | --- | --- | --- | --- |
| 1 | *INS1BP* | 4 | F | a |
| 2 | *INS1BP* | 4 | F | a |
| 3 | *INS1BP* | 4 | F | a |
| 4 | *INS1BP* | 4 | F | a |
| 5 | *INS1BP* | 6 | F | b |
| 6 | *INS1BP* | 6 | F | b |
| 7 | *INS1BP* | 6 | F | b |
| 8 | *INS1BP* | 10 | F | c |
| 9 | *INS1BP* | 10 | F | c |
| 10 | *INS1BP* | 10 | F | d |
| 11 | *DEL12BP* | 4 | F | e |
| 12 | *DEL12BP* | 4 | F | e |
| 13 | *DEL12BP* | 4 | F | e |
| 14 | *DEL12BP* | 4 | F | e |
| 15 | *DEL12BP* | 6 | F | f |
| 16 | *DEL12BP* | 6 | F | f |
| 17 | *DEL12BP* | 6 | F | f |
| 18 | *DEL12BP* | 6 | F | f |
| 19 | *DEL12BP* | 6 | F | f |
| 20 | *DEL12BP* | 6 | F | g |
| 21 | *DEL12BP* | 6 | F | g |
| 22 | *DEL12BP* | 6 | F | h |
| 23 | *DEL12BP* | 6 | F | h |
| 24 | *DEL12BP* | 10 | F | i |
| 25 | *DEL12BP* | 10 | F | i |
| 26 | *DEL12BP* | 10 | F | i |
| 27 | Wild-type NOD | 4 | F | j |
| 28 | Wild-type NOD | 4 | F | j |
| 29 | Wild-type NOD | 4 | F | j |
| 30 | Wild-type NOD | 4 | F | j |
| 31 | Wild-type NOD | 4 | F | j |
| 32 | Wild-type NOD | 6 | F | k |
| 33 | Wild-type NOD | 6 | F | k |
| 34 | Wild-type NOD | 6 | F | k |
| 35 | Wild-type NOD | 6 | F | k |
| 36 | Wild-type NOD | 6 | F | l |
| 37 | Wild-type NOD | 6 | F | m |
| 38 | Wild-type NOD | 6 | F | l |
| 39 | Wild-type NOD | 6 | F | n |
| 40 | Wild-type NOD | 6 | F | n |
| 41 | Wild-type NOD | 6 | F | m |
| 42 | Wild-type NOD | 10 | F | j |
| 43 | Wild-type NOD | 10 | F | j |
| 44 | Wild-type NOD | 10 | F | j |

**Supplementary Table S8 (Excel document Tab1) – Detected microbial species**

**Supplementary Table S9 (Excel document Tab2) – Robustly detected species**

**Supplementary Table S10 (Excel document Tab3) – Pathway set enrichment (microbiome)**

## Materials and Methods

### Mice

NOD/ShiLtJ mice were purchased from Charles River (Italy) for the purpose of generating the *Dexi*-disrupted mice used for this project. They were bred and maintained under a Home Office License and with the approval of the Local Animal Welfare Ethics Review Board (Clinical Medicine, University of Oxford). Mice were housed in same sex littermate groups of up to 6 mice in individually ventilated cages and were fed a standardized autoclaved commercial diet *ad libitum.* Standard conditions in the facility include shredded paper bedding and a 12 hour light-dark cycle. All animals were inspected daily and weekly urine testing for diabetes was undertaken from the age of 9-10 weeks. Mice were sacrificed as soon as glycosuria was detected, which was the endpoint of the cumulative diabetes incidence study.

### Generation and breeding of Dexi-disrupted NOD mice by CRISPR genome editing in NOD oocytes

A target site for CRISPR/Cas9 mutagenesis directed against the first exon of *Dexi*, [5’-CGTCCAGGTGGGCTGCGACC-3’] which contains the entire coding sequence of the protein, was designed using the CRISPOR algorithm (*71*). This target site was predicted by the algorithm to be genome-unique and in fact, a match to only a single putative off-target site (an intergenic region on a different chromosome) was only theoretically achievable with hybridization mismatches at 3 nucleotide positions within the target site, with also no predicted off-target sites predicted when 4 nucleotides were mismatched. As such, the target site for CRISPR/Cas9 mutagenesis selected for the *Dexi* loss-of-function has a high specificity score of 90 (*72*). The sgRNA was synthesized by *in vitro* transcription of a synthetic DNA template by T7 polymerase using the MEGAshortscript™ T7 Transcription Kit (ThermoFisher Scientific), purified using the MEGAclear Kit (ThermoFisher Scientific) and diluted prior to microinjection in 10 mM Tris.HCl pH7.5, 0.1 mM EDTA pH8.0. Female NOD mice at 6 weeks of age were superovulated and mated with male NOD mice. Fertilized zygotes were harvested from plugged females and microinjected with ribonucleoprotein preparation of sgRNA (at 20 ng/μl) complexed with 100 ng/ul Cas9 protein (Caribou Biosciences, Inc). Zygotes were cultured overnight and 47 of the resulting 2-cell embryos were implanted into 3 pseudopregnant foster CD1 females and carried to term. 19 live pups were obtained and genotyped for the presence of disruptive mutations at the CRISPR/Cas9 target site by PCR followed by Sanger sequencing.

Genotyping of pups was performed following PCR with Q5 HotStart Hi-fidelity DNA polymerase enzyme (New England Biolabs, Ipswich, MA, USA) using Sanger sequencing with primers directed at the first exon of *Dexi* (**Supplementary Table S2**). Mice with *Dexi* mutations were back-crossed onto the NOD background and resulting pups genotyped by Sanger sequencing to confirm the nature of the individual mutations. From this point forwards genotyping was undertaken by PCR and restriction fragment length polymorphism analysis after 6 hour digestion with *AvaII* (New England Biolabs, USA) for the *INS1BP* 1bp insertion mutation (**Supplementary Figure 2A**) or by PCR alone followed by 4% agarose gel electrophoresis for the *DEL12BP* 12bp deletion mutation (**Supplementary Figure 2B**).

Once it had been established that homozygous *Dexi* disruption was not lethal by evaluating offspring of heterozygous intercrosses, *INS1BP* and *DEL12BP* homozygous *Dexi*-disrupted mice were bred using homozygote intercrosses for the remainder of the project. Control NOD mice were generated by intercrossing wild-type NOD mice resulting from the original heterozygous intercrosses and these mice were used to breed control mice for subsequent experiments. Pups were weaned into same-sex littermate cages at 3 weeks of age. Mice were allocated to experimental groups according to genotype, age and sex. Randomization was not required since no additional interventions were undertaken. Wherever possible, data were collected and analyzed in a blinded fashion, as indicated in methods for individual experiments.

### Western blotting

SDS-PAGE gel electrophoresis was undertaken using 4-20% pre-cast Tris-Glycine gels (NuSep, Homebush, Australia) and PVDF blotting membrane. Tissue lysates from age matched nondiabetic female mice were prepared in RIPA buffer (with c0mplete Protease Inhibitor Cocktail – Sigma Aldrich, Gillingham UK) and 50ug total protein per well was loaded. Positive control Dexi protein lysate was obtained from Novus Biologicals (NBL1-09841, Abingdon, UK). Skimmed milk (5%) was used as a blocking agent in 0.5% Tween Phosphate Buffered Saline. Anti-DEXI antibodies were obtained from Proteintech (14811-1-AP, Rosemont, IL, USA), Sigma Aldrich (HPA041511, Gillingham UK) and Abgent (WA-AP 16938b, Hamburg, Germany). The secondary antibody was obtained from VectorLabs (Peterborough, UK – Goat anti-rabbit HRP conjugate, PI-1000) and the blots were developed with ECL Advance chemiluminescent reagent (Amersham, UK).

### Quantitative PCR

Tissues from 10-week old non-diabetic female mice were harvested immediately after sacrifice by cervical dislocation, and placed into Trizol for storage at −80 degrees. RNA was extracted using phenol-chloroform extraction and purified using Qiagen RNeasy columns, with an on-column DNAse digestion step. One microgram of RNA from each sample was used for cDNA synthesis with Bio Script enzyme and cDNA was diluted to 1:10 with purified water prior to use in qPCR reactions. Standard Master Mix (Primer Design, Southampton) was used for all reactions, which were undertaken using a Bio-Rad thermal cycler. Double-dye hydrolysis qPCR was used (TaqMan style) in duplex assays with beta actin (VIC-labelled probe) as the housekeeping gene. 5’ and 3’ Dexi assays (shown below) were designed and manufactured by PrimerDesign (FAM labelled probe), with the amplicons detected outlined on the image below. All samples were tested in triplicate and data shown for each mouse show mean +SD. Results are representative of multiple experiments.

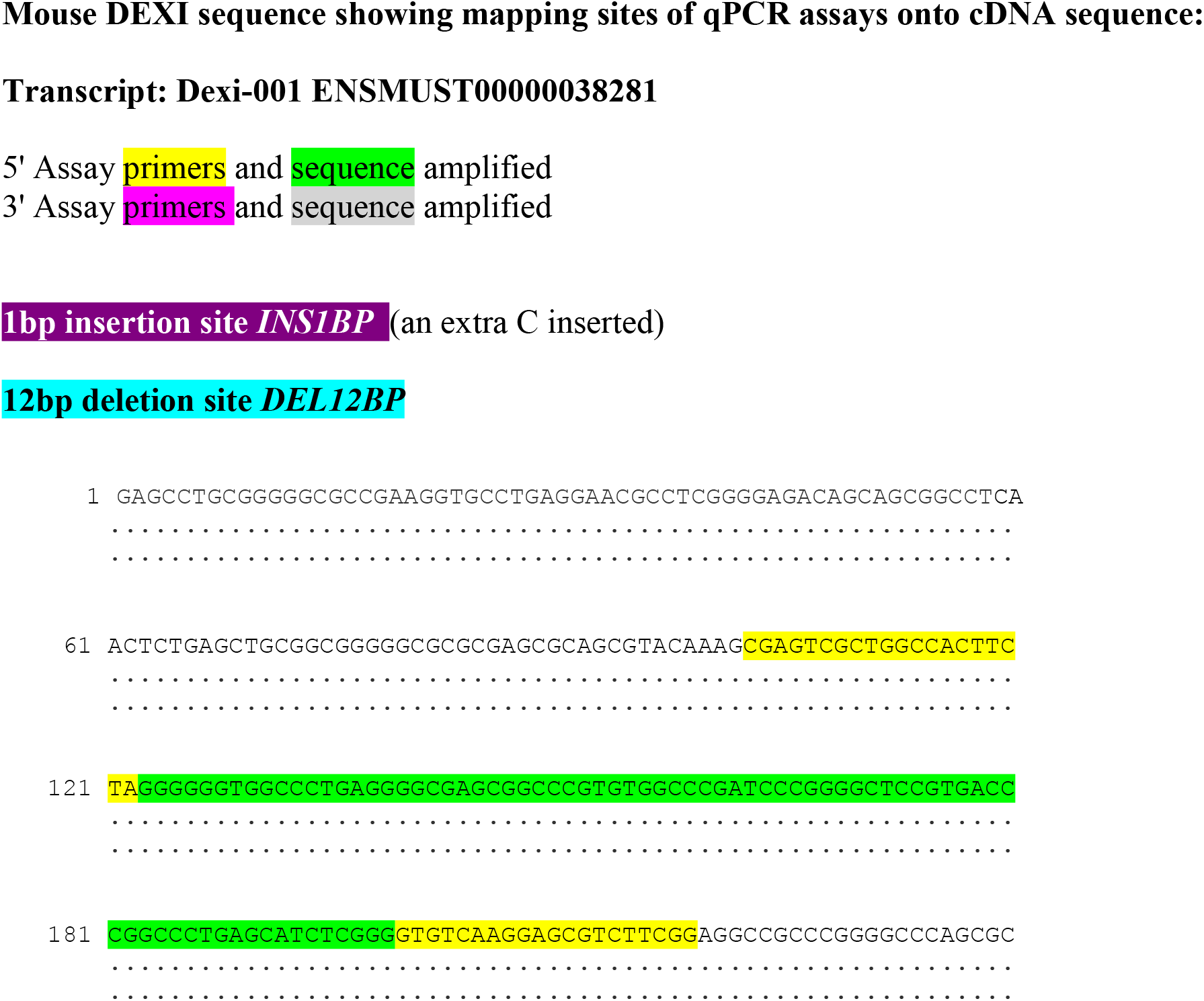
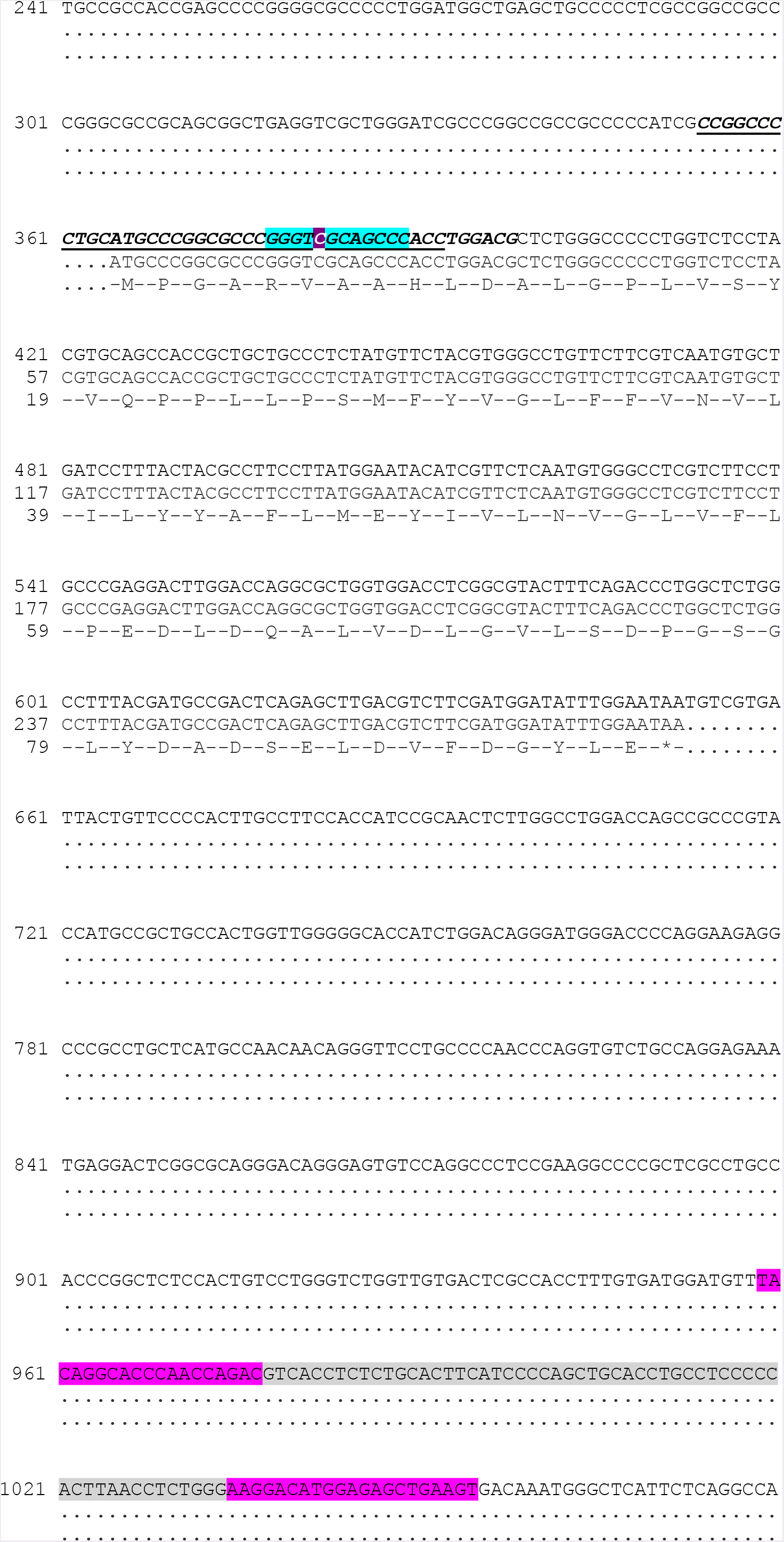

***Cumulative diabetes incidence study:*** From the age of 9-10 weeks, all mice in the colony underwent regular urine testing (weekly) for diabetes and were sacrificed as soon as glycosuria was detected. Since mice were kept in groups and cages labelled by genotype it was not possible for blinding to be undertaken with respect to genotype whilst urine testing was being undertaken. To avoid confounding hormonal effects, mice in breeding pairs and trios were not included in phenotyping studies or evaluation of diabetes incidence, although they received the same monitoring as non-breeding mice and were sacrificed as soon as glycosuria was detected. The study endpoint for diabetes incidence was set at 300 days. The ‘survival’ R package (version 2.41-3) was used to calculate diabetes-free survival, and the ‘survminer’ R package (version 0.4.0) was used to generate Kaplan-Meier plots.

***Histopathology (Insulitis study):*** Pancreas tissue from thirty three 6-week old female mice (**Supplementary Table S3**), was fixed for 24 hours in formalin before being transferred to 70% ethanol and stored at 4°C. Samples tissues were then embedded in paraffin and tissue sections cut in two planes. Following hematoxylin and eosin staining, an insulitis score (0 to 5) was evaluated by a board-certified veterinary pathologist in a blinded manner staining by evaluation of multiple islets in both planes for each mouse.

***Histopathology (Early phenotyping of first generations of Dexi-disrupted mice):*** Six littermate wild-type NOD and homozygous *Dexi*-disrupted NOD mice were sacrificed at 4-6 weeks of age and multiple tissues including pancreas, salivary gland, liver, spleen, kidney, bone marrow, thymus, lung, heart, gastro-intestinal tract and brain fixed in formalin for a minimum of 24 hours before embedding in paraffin. Sections were stained with hematoxylin and eosin before histopathological examination.

### Flow cytometry

After culling by cervical dislocation, spleen, thymus and femoral bone marrow (BM) from age-matched wild-type and *Dexi-*disrupted NOD mice were harvested into iced RPMI buffer with 10% fetal calf serum. A single cell suspension was made for each tissue and cells passed through a cell strainer before washing twice with ice-cold phosphate buffered saline/ 1% bovine serum albumin. After a red blood cell lysis step, cells were washed and re-suspended at 10 million cells / ml. One million cells per well were used for flow cytometry staining, which was performed on ice. Cells were stained with lineage-specific panels (**Supplementary Table S4**), with dead cells being excluded by Zombie Aqua staining (Biolegend, SanDiego, CA, USA). A mouse FC blocker (TruStain fcX™ (anti-mouse CD16/32) Biolegend) was included in the blocking step. For thymus, only the T cell panel was used. Data were acquired on a FACSCANTO machine and analyzed using FlowJo software. Fluorescence minus one controls and single-stained compensation controls were used to ensure that gates were set correctly and that fluorescently labelled antibodies in panels were compatible. For each subset identified, percentage of each subset within the population was calculated and this was converted to absolute number by multiplication by the total cell count from that tissue. Representative data from 4 mice (2 wild-type NOD and 2 *Dexi*-disrupted mice) are shown, and the experiment was repeated several times with similar results.

#### Flow cytometry gating strategies were as follows

Live B cells in the bone marrow were classified by Hardy fraction. B220+CD43+ cells were classified into Fraction A : early uncommitted cells BPI-CD24lo; Fraction B : rearranged H chain cells BPI-CD24hi; Fraction C : B cells post-H chain rearrangement BPI+CD24hi. B220+CD43-cells were classified into Fraction D : B cells rearranging L chain: IgM-IgD-; Fraction E: immature B cells with rearrangements finished IgM+IgD-; Fraction F: mature B cells IgM+IgD+. Live peripheral B cells in the spleen were classified as B cells according to B220+CD19+ status and then sub-classified as follicular (CD21+CD23+), marginal zone (CD21+CD23-) and transitional B cells (CD21-CD23-).

Live myeloid cells in the bone marrow and spleen were identified by their CD3-B220-status initially and subclassified into neutrophils (Gr-1+CD11b+) and MonoMac cells (Gr-1-CD11b+). MonoMac cells were subsequently further subclassified into monocytes (Ly6c+F4/80+); macrophages (Ly6c-F4/80+) and eosinophils (Ly6c-F4/80-).

Live T cells in the spleen were identified by their CD4+ and CD8+ status initially. CD4+ cells were subclassified into Tregs (CD62+CD25+); CD4+ and CD8+ T lymphocytes were also subclassified into effector cells (CD62-CD44+), memory cells (CD62+CD44+) and naïve cells (CD62+CD44-).

Live T cells in the thymus were sub-classified by CD4 and CD8 status as double negative (DN), double positive (DP), CD4+CD8- or CD4-CD8+. Double negative cells were further sub-classified by developmental stage into Q1 (CD44+CD25-), Q2 (CD44+CD25+), Q3 (CD44-CD25+) and Q4 (CD44-CD25-).

***Mice for IgA, IgM and IgG ELISA:*** The descriptive data for mice included in ELISA experiments are shown in **Supplementary Table S5**. Quantification of serum IgA, IgM and IgG was undertaken in age-matched young non-diabetic female mice using commercially available mouse ELISA kits, each containing a standard curve (ThermoFisher Scientific). Samples were tested in duplicate at a range of dilutions. Data were analyzed and displayed using GraphPad Prism. Since data were not normally distributed, wild-type NOD and *Dexi*-disrupted mice were compared (by group) using non-parametric testing (Mann-Whitney test), with significance set at p<0.05 and a Bonferroni correction for multiple testing. Results are shown from a single experiment but are representative of multiple experiments and figures were prepared using GraphPad Prism 7.

***Mice for IAA measurement:*** Aliquots of the same 36 mouse serum samples used for ELISA were shipped to the Barbara Davis Center for Childhood Diabetes, USA for measurement of IAA by radio-immunoassay. Investigators at this laboratory were blinded as to the genotype of each mouse.

***Metabolomics / lipidomics study mice:*** Five littermate female mice from two separate litters of each genotype (wild-type NOD, *INS1BP* and *DEL12BP*) were sacrificed by cervical dislocation at 6 weeks of age. Blood was immediately harvested into an EDTA tube by cardiac puncture and snap-frozen in a dry ice / isopropanol bath. Liver and spleen tissue were similarly snap-frozen immediately in separate tubes and all frozen samples stored at −80 degrees until shipping to the service provider (Metabolon, USA) on dry ice. One litter of each genotype was used for metabolomics studies (Metabolon HD4 platform) and one litter for lipidomics (Metabolon Complex lipids platform) studies. For the Short Chain Fatty Acid measurement experiment, 6-week old wild-type NOD (n=4), INSP1BP (n=3) and DEL12BP (n=3) mice from at least 2 litters per genotype were sacrificed by cervical dislocation at 6 weeks of age. Liver was immediately snap frozen as previously described. Blood was immediately harvested into a plain tube by cardiac puncture and after being allowed to clot for 10 minutes, was centrifuged and serum snap-frozen in a dry ice / isopropanol bath.

***Metabolomics sample processing for Ultra-high Performance Liquid Chromatography-Tandem Mass Spectroscopy (UPLC-MS/MS):*** Samples were processed according to Metabolon standard procedures and handled using the automated MicroLab STAR^®^ system from Hamilton Company. Several recovery standards were added prior to the first step in the extraction process for Quality Control purposes. To remove protein, dissociate small molecules bound to protein or trapped in the precipitated protein matrix, and to recover chemically diverse metabolites, proteins were precipitated with methanol under vigorous shaking for 2 min followed by centrifugation. The resulting extract was divided into five fractions: two for analysis by two separate reverse phase (RP)/UPLC-MS/MS methods with positive ion mode electrospray ionization (ESI), one for analysis by RP/UPLC-MS/MS with negative ion mode ESI, one for analysis by HILIC/UPLC-MS/MS with negative ion mode ESI, and one sample was reserved for backup. Samples were placed briefly on a TurboVap^®^ (Zymark) to remove the organic solvent. The sample extracts were stored overnight under nitrogen before preparation for analysis.

A pooled matrix sample, generated by taking a small volume of each experimental sample, served as a technical replicate. Extracted water samples served as process blanks. Metabolon QC standards were spiked into every analyzed sample. These steps allowed instrument performance monitoring and aided chromatographic alignment. Instrument variability was determined by calculating the median relative standard deviation (RSD) for the internal standards that were added to each sample prior to injection into the mass spectrometers. Overall process variability was determined by calculating the median RSD for all endogenous metabolites (i.e., non-instrument standards) present in 100% of the matrix samples, which are technical replicates of pooled samples.

All methods utilized a Waters ACQUITY ultra-performance liquid chromatography (UPLC) and a Thermo Scientific Q-Exactive high resolution/accurate mass spectrometer interfaced with a heated electrospray ionization (HESI-II) source and Orbitrap mass analyzer operated at 35,000 mass resolution. The sample extract was dried then reconstituted in solvents compatible to each of the four methods. Each reconstitution solvent contained a series of standards at fixed concentrations to ensure injection and chromatographic consistency. One aliquot was analyzed using acidic positive ion conditions, chromatographically optimized for more hydrophilic compounds. In this method, the extract was gradient eluted from a C18 column (Waters UPLC BEH C18-2.1×100 mm, 1.7 μm) using water and methanol, containing 0.05% perfluoropentanoic acid (PFPA) and 0.1% formic acid (FA). Another aliquot was also analyzed using acidic positive ion conditions; however it was chromatographically optimized for more hydrophobic compounds. In this method, the extract was gradient eluted from the same afore mentioned C18 column using methanol, acetonitrile, water, 0.05% PFPA and 0.01% FA and was operated at an overall higher organic content. Another aliquot was analyzed using basic negative ion optimized conditions using a separate dedicated C18 column. The basic extracts were gradient eluted from the column using methanol and water, however with 6.5mM Ammonium Bicarbonate at pH 8. The fourth aliquot was analyzed via negative ionization following elution from a HILIC column (Waters UPLC BEH Amide 2.1×150 mm, 1.7 μm) using a gradient consisting of water and acetonitrile with 10mM Ammonium Formate, pH 10.8. The MS analysis alternated between MS and data-dependent MS^n^ scans using dynamic exclusion. The scan range varied slighted between methods but covered 70-1000 m/z. Raw data files are archived and extracted as described below.

Data Extraction and Compound Identification : Raw data was extracted, peak-identified and QC processed using Metabolon’s hardware and software. Compounds were identified by comparison to library entries of purified standards or recurrent unknown entities. Biochemical identifications were based on three criteria: retention index within a narrow RI window of the proposed identification, accurate mass match to the library +/- 10 ppm, and the MS/MS forward and reverse scores between the experimental data and authentic standards. The MS/MS scores were based on a comparison of the ions present in the experimental spectrum to the ions present in the library spectrum.

***Metabolomics sample processing and data acquisition for the Complex Lipids Platform:*** For the EDTA blood samples, lipids were extracted from the blood in the presence of deuterated internal standards using an automated BUME extraction according to the method of Lofgren et al. (J Lipid Res 2012;53(8):1690-700). For tissue samples, lipids were extracted from samples via a modified Bligh-Dyer extraction using methanol/water/dichloromethane in the presence of deuterated internal standards. The extracts were dried under nitrogen and reconstituted in ammonium acetate dichloromethane : methanol. The extracts were transferred to vials for infusion-MS analysis, performed on a Shimadzu LC with nano PEEK tubing and the Sciex SelexIon-5500 QTRAP. The samples were analyzed via both positive and negative mode electrospray. The 5500 QTRAP was operated in MRM mode with a total of more than 1,100 MRMs. Individual lipid species were quantified by taking the ratio of the signal intensity of each target compound to that of its assigned internal standard, then multiplying by the concentration of internal standard added to the sample. Lipid class concentrations were calculated from the sum of all molecular species within a class, and fatty acid compositions were determined by calculating the proportion of each class comprised by individual fatty acids.

Short Chain Fatty Acids were measured by Mass Spectrometry (*73*) by Dr Zhanru Yu at the Target Development Institute, University of Oxford. Figures were prepared using GraphPad Prism 7.

### Bioinformatics and statistical analysis

***Metabolon analysis:*** Raw counts were rescaled for each biochemical to set the median equal to 1. Missing values were imputed with the minimum observed value for each compound. ANOVA contrasts were used to identify biochemicals that differed significantly between experimental groups. An estimate of the false discovery rate (*q*-value) was calculated to correct for multiple testing. The intersection of biochemical compounds with q < 0.05 and consistent fold change direction in both *Dexi-*disrupted lines versus wild-type NOD mice was taken to pinpoint biochemical compounds for further investigation. Metabolon pathway annotation for chemical compounds was used to identify pathways of interest. Bar and violin plots were generated in R (version 3.4.1). Pathway graphics were generated in Cytoscape using the Metabolync plugin.

For the SCFA measurement experiment, values were compared between groups using the Mann-Whitney test and a Bonferroni correction made for multiple testing.

***Fecal microbiome study:*** Fecal samples were collected non-invasively routinely from non-diabetic mice (either voided on handling or immediately after culling by cervical dislocation) and stored at −80 degrees for a maximum of 6 months until processed. Female mice were selected for this part of the study since the diabetes-acceleration phenotype had been detected in females. In addition, microbiome-dependent diabetes phenotypes in NOD mice are more apparent in females than males (*28, 29, 46, 74*). Samples were selected from the frozen archive based on age, sex and genotype, to ensure a wide variety of litters was represented. They comprised females of 4 weeks of age (n=13), 6 weeks of age (n= 22) and 10 weeks of age (n= 9). All three genotypes of interest were represented at all timepoints: total wild-type NOD (n=18), total *INS1BP* (n=10) and total *DEL12BP* (n=16), across a total of 12 different litters housed in single-sex separate cages of the same genotype (**Supplementary Table S7**). DNA was extracted from one fecal pellet per mouse with a combination of heat, chemical and bead dissociation using the Pure-Link Microbiome DNA purification kit (Thermo-Fisher, Waltham, US). DNA quality and purity was assessed by Nanodrop, to confirm a 260/280 ratio between 1.8-2.0 and 260/230 ratio of greater than 1.5. Fifty nanograms of fecally-derived DNA from each mouse was submitted to the High Throughput Genomics Service at the Oxford Genomics Centre, Wellcome Centre for Human Genetics. Forty four samples were tested in a blinded fashion across two Axiom Microbiome arrays (ThermoFisher, Waltham, US), with the remaining two spaces on each 24-sample array taken up with positive and negative controls.

Raw. CEL output files were analyzed using Axiom Microbial Detection Analysis Software (MiDAS), which makes species ‘Detected Target’ calls based on the Composite Likelihood Maximization (CLiMax) algorithm. The ‘pheatmap’ R package (version 1.0.8) was used to cluster mouse samples by euclidean distance and generate a heatmap of Detected Target microbial species. The ‘VennDiagram’ R package (version 1.6.17) was used to generate Venn diagrams of Detected Targets. To minimize the risk of batch artifacts, analyses were restricted to only Detected Target species found in both array runs and detected at each time point (4, 6, & 10 weeks of age), which also corresponded to species detected in more than one mouse line. Fisher’s Exact was used to test differences in the representation of these robustly detected microbial species in *Dexi-*disrupted versus wild-type NOD mice.

## Author contributions

Conceived and designed the study: LJD, CAO’C, JAT

Designed and generated the *Dexi*-disrupted mice: CP, BD

Undertook experiments: LJD

Undertook blinded histopathological analysis of tissues and immunohistochemistry: KH

Analyzed data: LJD, MDW, CO’C

Wrote the manuscript: LJD, CAO’C

Prepared figures: LJD, MDW, KH

Reviewed data, reviewed and edited the manuscript: JAT, CP, BD

## Ethics

All animal studies were conducted with ethical approval from the John Radcliffe Hospital Clinical Medicine Animal Welfare and Ethical Review Board and in accordance with the UK Home Office regulations (Guidance on the Operation of Animals, Scientific Procedures Act, 1986), under Project License PPL 30/3029.

## Funding

This work was supported by a Wellcome Trust Veterinary Postdoctoral Fellowship to LJD (WT096398MA) and by a pilot study grant from Diabetes UK to LJD and CO’C (15/0005171). MDW’s salary is supported by an EFSD/JDRF Lilly pilot grant (EFSD/JDRF/Lilly Programme 2016) to LJD and CO’C. LJD is currently supported by an MRC Clinician Scientist Fellowship (MR/R007977/1).

The work was supported by the Juvenile Diabetes Research Fund [5-SRA-2015-130-A-N], [4-SRA-2017-473-A-N]; the Wellcome Trust [107212/Z/15/Z]; National Institute of Health Research Oxford Biomedical Research Centre. JAT was in receipt of a National Institute of Health Research Senior Investigator Award.

No funding bodies had any role in study design, data collection and analysis, decision to publish, or preparation of the manuscript.

## Acknowledgments

We thank the High-Throughput Genomics Group and the transgenic core at the Wellcome Centre for Human Genetics (funded by Wellcome Trust grant reference 203141/Z/16/Z) for the production of the *Dexi* mutant mice and the generation of the microbiome array data. We thank Dr. Joy Archer FRCPath at the University of Cambridge for undertaking the initial histopathology work on the mouse tissues. We thank the staff of the Functional Genomics Facility, University of Oxford for their technical expertise in caring for the mice. We are grateful to Dr. Consuelo Anzilloti and Dr. Mukta Deobagkar for help and guidance with immunophenotyping the mice by flow cytometry. We thank Prof. Linda Wicker for valuable discussions and Dr Zhanru Yu for undertaking SCFA measurements.

